# Female site fidelity and repeated pairings across years in bull sharks (*Carcharhinus leucas*) inhabiting Fiji waters

**DOI:** 10.64898/2026.03.23.713736

**Authors:** Kerstin Glaus, Laura Benestan, Juerg Brunnschweiler, Floriaan Devloo-Delva, Sharon A. Appleyard, Ciro Rico

## Abstract

Understanding relatedness in sharks is challenging due to uncertainty in distributions, low population densities and difficulties in sampling across life stages. In Fiji, bull sharks (*Carcharhinus leucas*), with an effective population size estimate of ∼258, aggregate at the Shark Reef Marine Reserve (SRMR), but gravid females disperse at the end of the year to give birth in adjacent rivers. Questions remain regarding reproductive connectivity, female returns across years, and kinship structure. Using population genomics on 296 bull sharks across age classes (neonates, young-of-the-year, juveniles, and adults) collected over a decade at the SRMR and in three adjacent rivers, we assessed familial connections. Direct genetic links, including first- and second-degree relationships, connected SRMR adults with young age classes in the Navua and Rewa rivers, providing evidence of reproductive connectivity. Within rivers, genetic similarities across cohorts revealed reproductive philopatry. Remarkably, several individuals sampled years apart were assigned to the same sire-dam pairs, indicating repeated pairings across breeding seasons. However, the few related links detected between the SRMR and the rivers may reflect incomplete sampling. Altogether, bull shark reproduction in Fiji seems influenced by reproductive philopatry and repeated pairings, suggesting added complexity in their reproductive behaviour.

## 1. Introduction

Coastal shark populations are in marked decline, with cascading consequences for community structure, trophic interactions, and the stability of coastal ecosystems [1]. Once depleted, shark populations are slow to recover due to their life-histories, which generally include long lifespans, variable movement patterns, low fecundity, and late maturity [2–4]. Understanding their reproductive behaviour, population connectivity, and demography is therefore critical for conservation management [5, 6]. However, obtaining sufficient data across life stages and habitats remains challenging because of wide ranges and low population densities [7].

Genetic relatedness analyses are an important tool to investigate connectivity, philopatry, kinship, and effective population size. For example, in bull sharks (*Carcharhinus leucas*) from the south-eastern United States, high relatedness revealed by microsatellite data suggested limited genetic diversity [8]. Long-term monitoring of lemon sharks (*Negaprion brevirostris*) at Bimini showed that females return to the same nursery grounds every two years to give birth, a clear indication for reproductive philopatry [9, 10] and uncovered natal returns [9]. In rare species such as the speartooth shark (*Glyphis glyphis*), close-kin mark-recapture has been used to estimate adult abundance and connectivity between rivers, again revealing reproductive philopatry [11]. Relatedness has also been applied to explore social structure in lemon sharks [12] and the bluntnose sixgill shark (*Hexanchus griseus*) [13]. Collectively, genetic relatedness analyses can help to identify critical habitats and uncover behaviours that otherwise remain elusive [14].

The bull shark is a large predator with a circumtropical distribution that inhabits a wide range of coastal habitats [15]. As an euryhaline shark, it plays a critical ecological role across freshwater, estuarine, and marine environments [16, 17], including mediating connectivity between marine and freshwater food webs [18]. Bull sharks are viviparous with a biennial reproductive cycle, producing litters of 1-14 pups (mean 6-10) after a gestation period of 10-11 months [19]. Young bull sharks undergo ontogenetic niche shifts [20, 21], spending their early years in rivers and estuaries as predator refuges [22] before moving into marine habitats. Gene flow is generally maintained across shallow coastal waters, which act as dispersal corridors, whereas large oceanic distances and historical land bridges represent barriers [23, 24]. Females exhibit reproductive philopatry, returning to the same areas to give birth [25, 26]. Bull sharks face multiple threats, including targeted, opportunistic, and incidental capture in commercial and small-scale fisheries, habitat degradation from coastal development, pollution, and changing environmental conditions [27–30]. The species is now classified as Vulnerable on the International Union for Conservation of Nature (IUCN) Red List of Threatened Species [31].

In Fiji, bull shark fishery, ecology, and behaviour have been studied for over two decades [32–34]. For instance, genomic research has revealed that Fiji’s bull shark population is genetically distinct [23, 35], while individuals of all life stages are captured in the country’s small-scale fishery [29]. Adult bull sharks are abundant at the Shark Reef Marine Reserve (SRMR), a no-take zone and provisioning site off Viti Levu [36]. Social studies at the SRMR have documented enduring associations among adults persisting over months to years, suggesting stable social bonds within the population [37]. Younger life stages, including neonates, young-of-the-year (YoY), and juveniles, occur in several adjacent rivers that provide essential habitats [38, 39]. Acoustic tracking has further shown that these rivers are critical for completing the bull shark life cycle [40]. However, reproductive links between SRMR adults and riverine pups, kinship structure, and effective population size (*N_e_*) estimates remain unresolved.

Using thousands of genome-wide Single Nucleotide Polymorphism (SNP) loci supports accurate determination of family relationships and identification of different types of relatives, such as full and half siblings (Attard et al., 2018). While microsatellites have long been the most common markers for kinship studies in wild populations [6], SNPs offer greater statistical power for estimating relatedness [41, 42]. Here, we genotyped adult bull sharks from the SRMR which were sampled over a decade, alongside cohorts of young age classes sampled from three adjacent rivers in Viti Levu over up to seven years. Our objectives were to (1) investigate kin structure between the SRMR and adjacent rivers, (2) examine kin structure within rivers over time, (3) assess kin structure among rivers, and (4) estimate N_e_. Together, our findings provide a finer resolution of bull shark population structure in Fiji and can inform habitat management at the local scale.

## 2. Materials and Methods

### 2.1. Study area, target species and sample collection

The study focused on the waters surrounding the south coast of Viti Levu, Fiji (Figure 1). A total of 353 bull sharks were sampled across three rivers – Navua, Rewa, and Sigatoka – and the SRMR. To obtain candidate parents, white muscle samples (0.5 cm³) from adult bull sharks were collected between 2007 and 2017 in the SRMR. Initially, a custom-made pole spear and later a small pneumatic speargun with a manually adjusted sharp tip and barbs were used to sample free-swimming bull sharks. A certified PADI Divemaster and marine biologist collected samples, averaging one shark per week or every fortnight. Throughout the sampling period, associated biological data were recorded, including sex and estimated age class whenever possible. Muscle samples were preserved in 90% ethanol.

**Figure 1.**
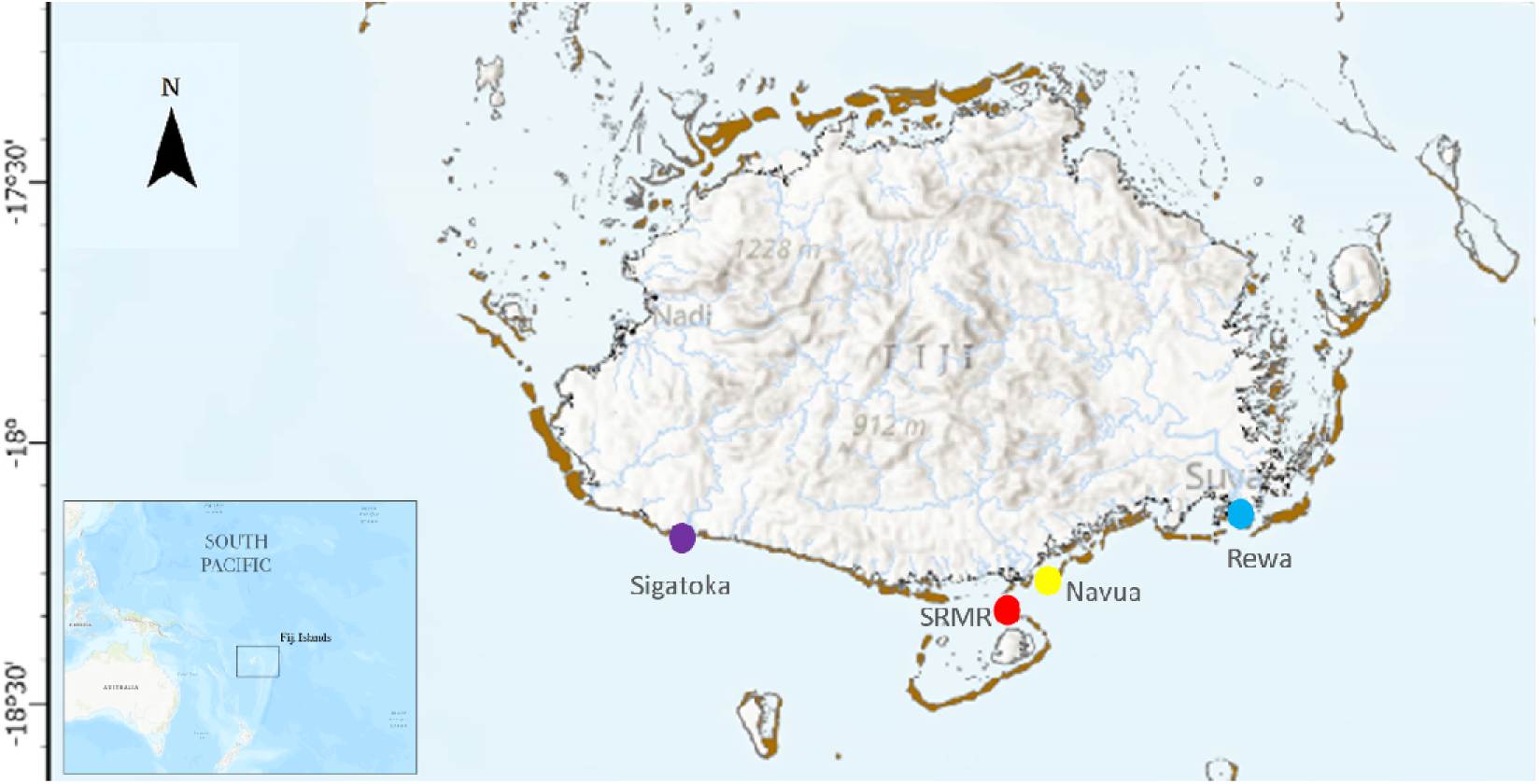
Map of Fiji’s main island, Viti Levu, indicating the four sampling locations from west to east: Sigatoka River, SRMR, Navua River, and Rewa River.

To obtain candidate offspring, non-destructive tissue sampling took place first year-round and then shortly after pupping (e.g., during austral summer) for young age classes of bull sharks in the Navua, Rewa, and Sigatoka rivers on the southern coast of Viti Levu (for details, see Glaus, Brunnschweiler [39]). In short, boat-based fisheries-independent surveys were conducted between 2014 and 2018. Surveys targeted the lower to mid reaches and estuaries, beginning at low tide and lasting 2–6 h per day. Sampling was conducted with a 150 × 3 m gillnet, which was inspected every 20–35 min to minimize risk of mortality due to asphyxia. Captured bull sharks were transferred to an onboard tank filled with river water, and the following parameters were recorded: total length (TL), umbilical scar condition (open, semi-healed, healed according to Duncan and Holland [43]). In addition, carcasses or body parts (e.g., fins) of bull sharks captured in the Rewa and Navua rivers were provided by known local small-scale fishers. We informed the fishers about the study’s purpose and encouraged them to release any live sharks. We did not provide any financial incentives in exchange for bull shark carcasses or body parts. We extracted a small fin clip, approximately 1 cm in length, from the free rear tip of the captured individuals. The fin clips were stored in Eppendorf tubes filled with 90% ethanol and kept at −20°C until shipment and further processing.

### 2.2 DArT-Seq^TM^ library preparation and sequencing

All bull shark tissue samples were sent to Diversity Arrays Technology (DArT™) in Canberra, Australia, where genomic DNA was extracted with a Macherey-Nagel NucleoMag 96 Tissue kit [44]. The DNA was processed for reduced representation library construction, sequenced, and genotyped using the DArT-Seq™ protocol, following a previously developed and tested complexity reduction protocol for scalloped hammerhead sharks [45]. Briefly, genome complexity was reduced using a double restriction digest with a PstI and SphI methylation-sensitive restriction enzyme combination. Libraries were sequenced on an Illumina HiSeq 2500 platform, and raw reads were processed with Illumina CASAVA v.1.8.2 software to assess of read quality and sequence representation.

### 2.3 SNPs filtering – quality control

We genotyped 26,015 SNPs in 353 individuals. We examined heterozygosity patterns and applied a filter with an upper threshold of 0.5 and no individuals were removed. Using the dartR package (Gruber et al., 2018), we kept SNPs that had more than 95% of data available, a minor allele frequency (MAF) over 5%, met the Hardy Weinberg Equilibrium (HWE) with a significant p-value of 0.05, and had a linkage disequilibrium with R^2^ < 0.20. We kept individuals with less than 10% missing data (Table S1).

### 2.4 Pedigree reconstruction

To infer both full and half sibling assignments, we used the software program COLONY [46]. Runs in COLONY were performed across all samples, with the program using a group-wide maximum likelihood method to infer sibship. The likelihood-based algorithms in COLONY partitioned the sampled individuals into sibling groups to maximize the probability of the observed data. The higher the probability, the more confident of the COLONY calls on kin relationships. Here, we set a probability threshold of 0.8. Polygamous mating systems were assumed for both sexes to allow for the assignment of full and half siblings. The software also incorporates sibship prior information, using a rate of 1 (weak sibship prior), as indicated by multiple initial runs at various rates (weak, medium and strong sibship prior). For each data set, ten long runs with high precision, based on different random seeds, were conducted to confirm assignment reliability using a maximum likelihood configuration and a Markov chain Monte-Carlo (MCMC) algorithm. Relationships were inferred in COLONY from population allele frequencies, which were adjusted in small or uneven datasets by adding pseudocounts to avoid zero frequencies and biased parentage or sibship assignments. The number of full and half siblings identified by COLONY were visualized using the *circlize* R package [47].

To visualise family cluster genetic relationships and confirm the assignments observed in COLONY, we constructed networks based on the Wang relatedness estimator [48] using the *coancestry* function from the RELATED R package [49]. This estimator provides a measure of pairwise genetic relatedness (r) ranging from −1 to 1, where 0.5 corresponds to full siblings or parent-offspring, 0.25 to half siblings, values greater than 0.5 may indicate inbred full siblings or duplicates, and 0 indicates unrelated individuals (Figure S1). We applied the relatedness estimator to identify and filter out recaptured individuals, retaining only those with r>0.6. After this filtering step, our dataset was reduced to 296 unique individuals out of the 353 genotyped samples. Using the resulting relatedness matrix, we built networks by applying the *graph_from_data_frame* function from the IGRAPH package [50], allowing us to explore family patterns and highlight site fidelity and/or mate fidelity. Finally, we applied the *gl.filter.parent.offspring* function in dartR package to investigate parent-offspring relationship based on the frequency of inconsistent loci, defined as cases where the parents are homozygous for the reference allele while the offsprings is homozygous for the alternative allele.

### 2.5 Effective population size

We estimated the genetic effective population size (*N_e_*) for the bull shark population in Fiji using NeEstimator v.2.1 [51]. Estimates were not separated by sex due to the small sample size of male individuals. The analysis looked at all the data, including both related and unrelated sharks, using the linkage disequilibrium method and a random mating model to calculate estimates with 95% confidence intervals.

## 3. Results

### 3.1 Kinship structure

Following filtering and duplicates removal (see above), a total of 296 bull shark individuals originating from three rivers – Navua (*n* = 28), Rewa (*n* = 166), and Sigatoka (*n* = 9) – as well as from the SRMR (*n* = 93) were successfully genotyped at 4,146 SNPs and included in the dataset for pedigree reconstruction. Assignment analyses identified 375 related pairs (0.8%), comprising 146 full siblings and 229 half siblings (Table S2). Full-sibling pairs were primarily detected within individual locations: 12 pairs in the Navua River (0.03%), 81 in the Rewa River (0.2%), one in the Sigatoka River (2.29×10^−3^ %), and seven in the SRMR (0.1%). Additionally, related pairs were identified between the SRMR and each of the three rivers: eight pairs between the SRMR and the Navua River, 30 pairs between the SRMR and the Rewa River, and one pair between the SRMR and the Sigatoka River (Figure 2a, b).

**Figure 2.**
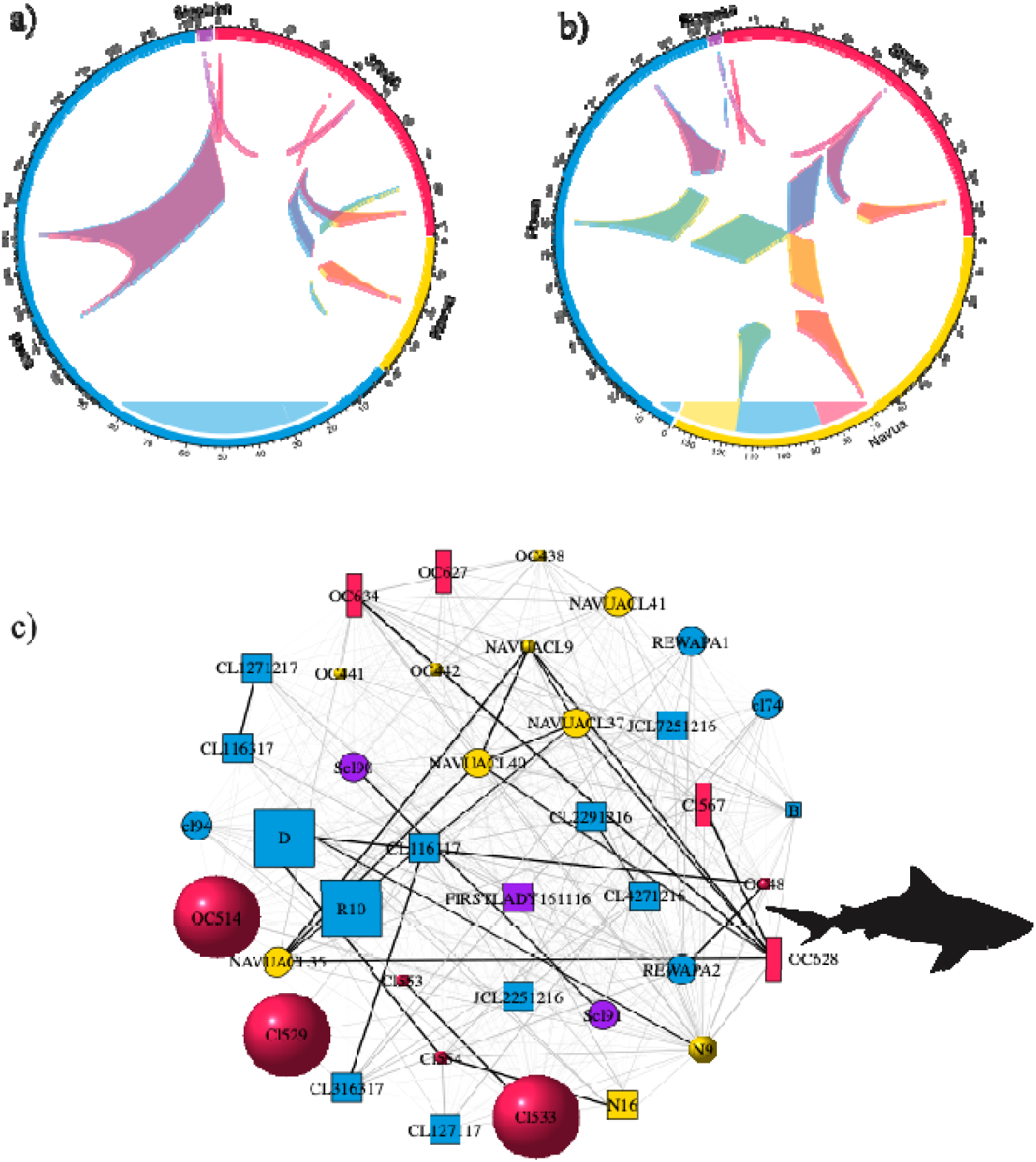
Connectivity across the SRMR and rivers. (a-b) Chord diagram illustrating the number of full-sibling (a) and half-sibling (b) relationships shared across sampling locations. (c) Network depicting kinship links within the largest genetic cluster (ClusterIndex 2) identified by COLONY, including individuals from all three rivers. First-degree (black lines) and second-degree (grey lines) relationships are shown. Each node represents a shark and is labelled with its individual ID. Node color corresponds to sampling locations, consistent with the sampling map. The node shape indicates the year of sampling; where no year was available, the node is small. Node size reflects the age class of the individual (neonate, young of the year, or adult).

A notable example of a first-degree relationship spanning rivers involves individual OC628, sampled in 2011 in the SRMR (likely at least subadult), which was identified as a full sibling to four individuals (including three neonates) from the Navua River: one sampled in 2014 (NavuaCl9) and three sampled in 2017 (NavuaCl35, NavuaCl37, NavuaCl40; Figure 2c). Similarly, 16 of the 30 full-sibling pairs identified between the SRMR and the Rewa River involved adult individuals sampled in the SRMR between 2008 and 2010 and neonates sampled in the Rewa River in 2016 and 2017 (Table S2). These findings indicate connectivity between the SRMR and these river systems over multiple years.

Second-degree kinship analyses further revealed strong genetic connections between the rivers and the SRMR, except for the Sigatoka River, where fewer bull sharks were detected despite sampling efforts comparable to those in the Rewa River over two consecutive years. The number of second-degree related pairs ranged from one pair (Sigatoka/SRMR or Sigatoka/Rewa) to 28 pairs (Rewa/SRMR). Again, neonates sampled in the Rewa and Navua rivers between 2016 and 2017 shared second-degree relationships (*n*=32) with adults sampled in the SRMR between 2007 and 2010, highlighting long-term intergenerational connectivity across these systems. Cross-cohort half-siblings indicate repeated reproduction by one parent across years, whereas same-cohort half-siblings may reflect dispersal from the natal site.

### 3.2 Reproductive Philopatry

When examining kin relationships within river systems over time, we identified several full- and half-sibling pairs sampled from YoY more than two years apart. A total of 15 individuals exhibited kinship links with others collected at least two years later, which coincides with the species’ biannual reproductive cycle. This indicates they belong to the same family but represent different cohorts (Table S2). In the Navua River, individual NavuaCl40, NavuaCl35, and NavuaCl37 were identified as full siblings of NavuaCl9, despite being sampled three years apart (Figure 2c, Figure 3a). In the Rewa River, YoY individuals R11, R15, R12, R16, and D sampled in 2015 were half siblings of neonates (Cl46, Cl60, Cl9, Cl36, Cl74, Cl78, and RewaPa2) sampled in 2017 (Figure 3b). In the SRMR, adult Cl530 sampled in 2010 was identified as a full sibling of 101016SRadultF, sampled in 2016. Similarly, SR17Blunt, sampled in 2017, was a full sibling of OC430 and OC630, sampled in 2010 and 2011, respectively (Figure 3c). This temporal continuity in kinship structure provides compelling evidence for reproductive philopatry in bull sharks.

**Figure 3.**
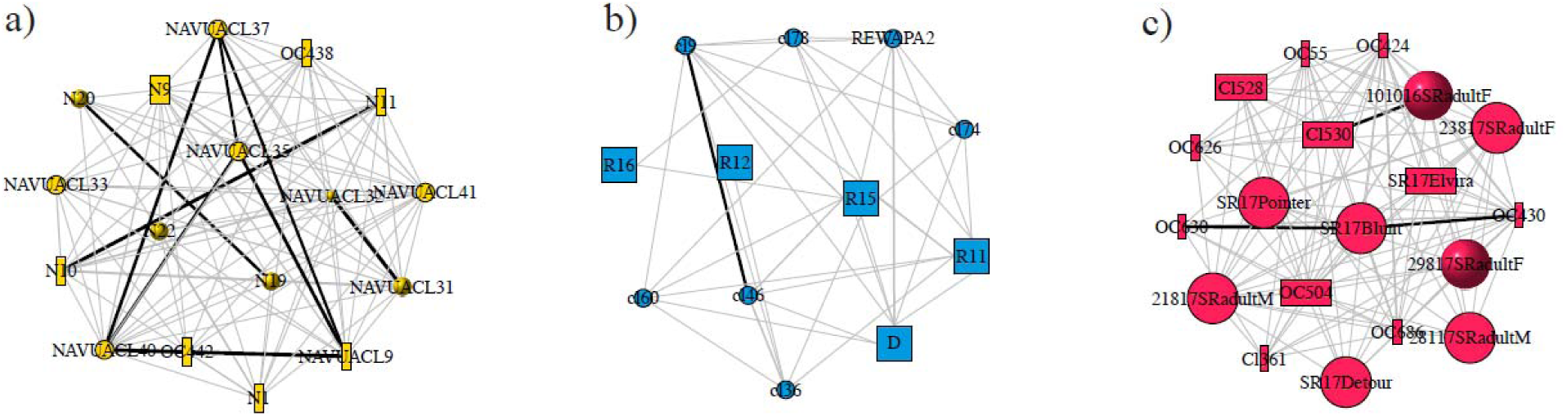
Site fidelity across the years within river systems and the SRMR. Networks of the kin relationships within a single-family cluster, illustrating evidence of multi-year site fidelity. First-degree (black lines) and second-degree (grey lines) relationships are displayed. Each node represents a shark, labelled by its individual ID. Node colour corresponds to sampling locations, consistent with the sampling map. The node shape reflects the year of sampling: rectangle (before 2014), square (2015), sphere (2016) and circle (2017). Nodes with no year information are shown as smaller points. Node size corresponds to the age class of the individual (neonate, young of the year, or adult).

### 3.3 Repeated pairings across years

Fifty-four sharks sampled across different years (with at least a two-year gap) were assigned to the same family clusters and shared both their mother and father, indicating repeated pairings across years (Table 1). These individuals belong to 13 family clusters, each exhibiting a consistent pattern: young sharks sampled after 2014 in either the Navua or Rewa rivers are genetically linked to individuals sampled earlier in the SRMR or to adults present during the same period. For instance, one cluster includes the adult individual SR17Elvira, sampled in the SRMR in 2017. This shark shares both a mother (ID 109) and a father (ID 110) with five neonates (Cl22, Cl23, Cl25, Cl53, and Cl75) sampled in the Rewa River in 2015 and 2017. Family cluster sizes range from two individuals (e.g., Cluster Index 5) to six individuals (e.g., Cluster Index 48), and the temporal span between related individuals can extend up to ten years. A notable example is Cluster 18, in which mother 25 and father 22 produced OC51 in 2007 and subsequently produced Cl52, sampled in the Rewa River in 2017, illustrating a decade-long reproductive continuity within the same parental pair.

**Table 1.**
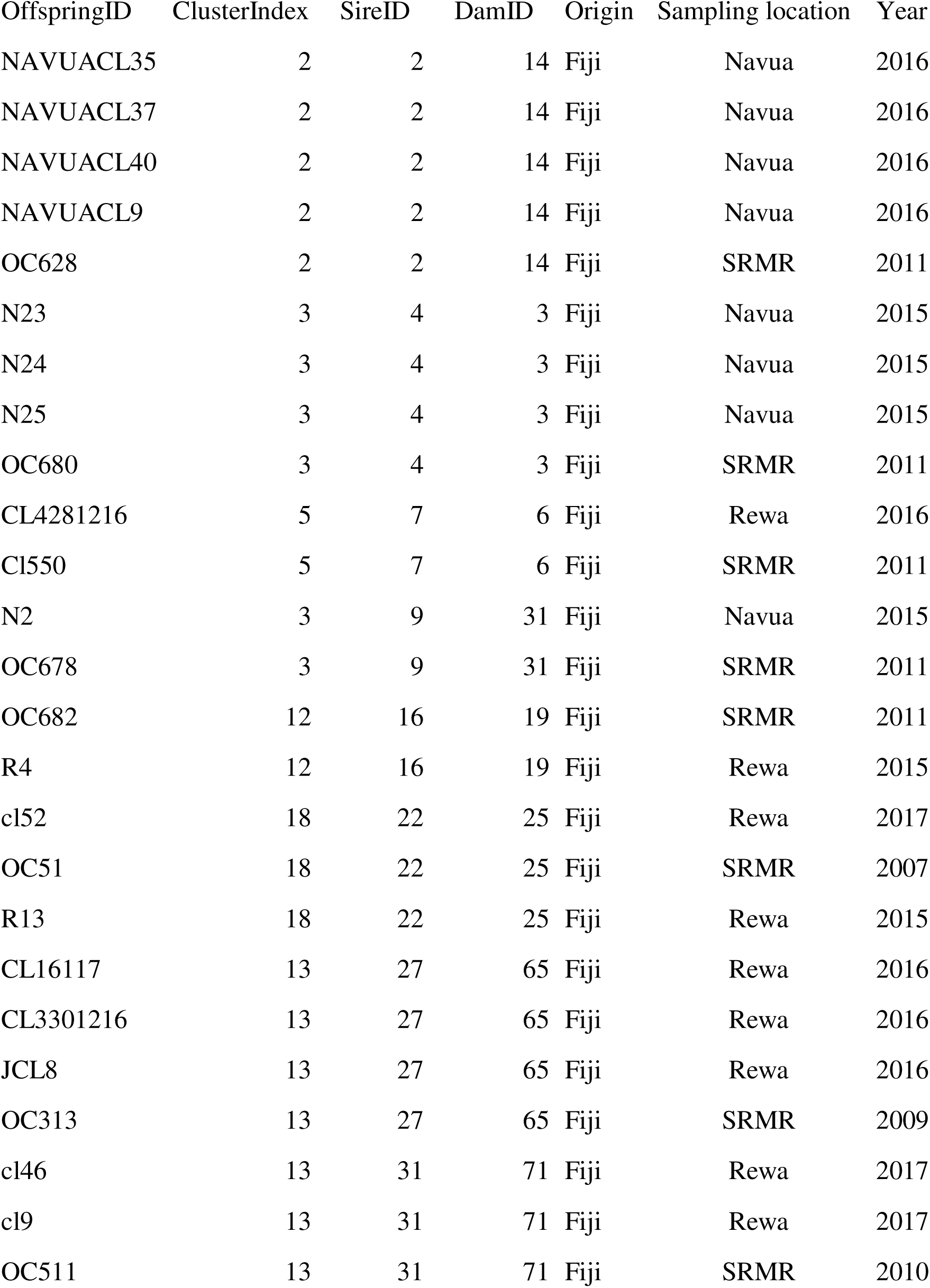
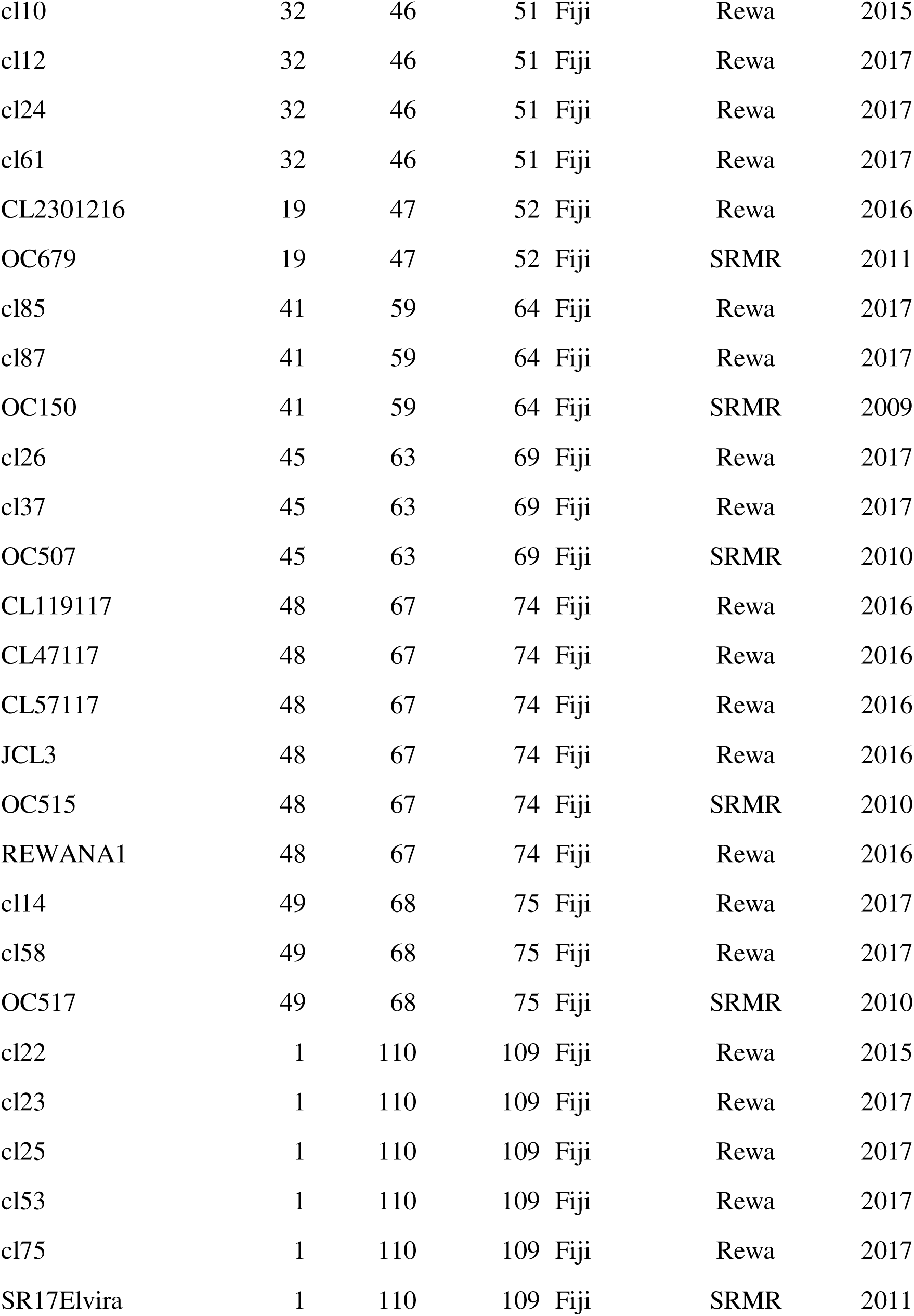
Repeated pairings, including cluster index, sire and dam IDs as identified by COLONY, and the origin, sampling location, and year of sampling.

### 3.4 Effective population size

The N_e_ estimates, including related individuals, amounted to 149.6, nearly doubling to 283.2 when related individuals were excluded. Both estimates were associated with narrow confidence intervals (149.3 − 149.9 and 281.8-284.6), indicating high precision around the estimates (Table 2).

**Table 2.** Estimates of effective population size (N_e_) for all individuals (n = 296) and for unrelated individuals (n = 193), with minor allele frequency >0.05, and lower and upper confidence intervals (CI low and CI high).

|  | **#samples** | **# of loci** | **$N_e$ (MAF $> 0.05$)** | **CI low** | **CI high** |
| --- | --- | --- | --- | --- | --- |
| All | 296 | 4,514 | 149.6 | 149.3 | 149.9 |
| Unrelated | 193 | 4,514 | 283.2 | 281.8 | 284.6 |

## 4. Discussion

Shark mating behaviour remains largely elusive due to the challenges of directly observing reproductive events in the wild. Consequently, genetic approaches such as parentage assignment and relatedness analyses have become key tools for uncovering reproductive dynamics [5, 7]. Here, we sampled candidate parents and offspring of bull sharks over more than a decade, spanning multiple cohorts, marine and estuarine locations and thousands of SNPs. This long-term genetic profiling provides new insights into the bull shark life cycle in Fiji. Previous ecological surveys documented neonate bull sharks in the sampled rivers during the austral summer parturition season [38, 39], and acoustic tracking has confirmed that gravid females from the SRMR enter the river estuaries [40]. While earlier work demonstrated the importance of these rivers as parturition sites, our genetic analyses revealed direct first and second-degree links between adult bull sharks from the SRMR and neonates, YoY, or juveniles in the Navua and Rewa rivers. These findings further highlight the critical role of riverine habitats in the bull shark life cycle and underscore the interconnectedness of marine and freshwater ecosystems in Fiji.

Site fidelity has been documented in several shark species [7, 52–55]. By returning to the same feeding grounds, mating grounds, or nurseries, sharks can repeatedly make use of predictable prey or habitat resources [56]. Reproductive philopatry further ensures that offspring are born in habitats that have historically supported survival and may help reinforce genetic adaptations to local conditions [53]. In this study, we also confirmed reproductive philopatry in bull sharks in Fiji, with full and half siblings detected across cohorts in the Navua and Rewa rivers, but not in the Sigatoka River. In the Sigatoka River, only one genetic link was detected, reflecting the relatively low number of bull sharks there contributing to the observed reproductive philopatry. The lower number of bull sharks in the Sigatoka River may be due to the shorter sampling period, despite considerable effort, combined with reduced suitability of the river due to environmental alterations and a reported decline in shark presence [29, 39]. Also, dispersal from natal sites warrants targeted study, as postnatal movement patterns of Fiji’s young age classes of bull sharks remain unknown.

Across the rivers, the low number of genetic links may indicate limited sampling or that individuals originate from outside the SRMR. Indeed, neonate bull sharks have been documented in the Dreketi River on Vanua Levu [57], the country’s second-largest island, and the species has also been reported in Tonga, approximately 800 km away [58]. Bull sharks have the capacity for long-distance movements and return migrations; notably, a pregnant female bull shark was tracked migrating more than 2,000 km from the Seychelles to southeast Madagascar and back [26]. Expanding sampling locations across Fiji and beyond could reveal additional relatedness, including parent–offspring, links.

Genetic studies examining relatedness in sharks face the practical challenge of requiring relatively larger sample sizes collected over extended time frames, a difficulty compounded by the long generation times of many species. Consequently, most relatedness studies to date have relied on a limited number of litters or siblings sampled within a single year [5] [but see 9]. In contrast, our decade-long dataset, spanning multiple cohorts of adults and young age classes, provides the first evidence of repeated pairings across years between the same male and female bull shark. We estimated the effective population size of bull sharks in Fiji at just above half of the commonly cited threshold of 500 [59], and well below the recommended 1,000 considered [60] sufficient to reduce inbreeding and maintain evolutionary potential in marine organisms. Most likely the small population size would increase the probability of repeated pairings under the random mating hypothesis [61]. Alternatively, such pairings could reflect behavioural strategies that confer indirect fitness benefits, for example by enhancing offspring viability through compatible or high-quality mates. For example, few larger males could dominate the access to the few females. In addition, female selectivity would favour the larger males [62]. Overall, this further reduces the pool of available partners, and increases the likelihood of repeated pairings. The findings may also result from physiological processes such as long-term sperm storage. In some elasmobranchs, females are known to store sperm for periods ranging from weeks to years [63, 64]. In bull sharks from La Réunion, Pirog, Magalon [19] inferred a sperm storage period of roughly 4–5 months. However, our study identified 13 family clusters involving repeated parental pairings, including cases of reproductive continuity over a span of up to ten years, going beyond the temporal viability of stored sperm.

Comparable evidence for repeated pairings in other shark species is scarce. In lemon sharks, Feldheim, Gruber [55] reconstructed parental genotypes across six breeding seasons at Bimini using 910 individuals (13 adults, 130 subadults, 222 juveniles, 532 neonates, and 13 of unknown age). Despite this large sample size, only a handful of repeated male–female pairings were detected, and these could not be clearly distinguished from sperm storage events. The difference may partly reflect species-specific strategies, but also differences in sampling design: whereas Feldheim’s dataset was dominated by juveniles and neonates, ours included proportionally more adults and spanned a broader temporal scale across both rivers and the SRMR. Whether these patterns reflect genuine repeated pair bonds or post-copulatory mechanisms such as sperm storage remains uncertain. Nevertheless, the decade-long continuity observed here suggests that repeated pairings may form part of bull shark reproductive biology in Fiji, with similar dynamics potentially present in other insular island populations [23]. Non-exclusive demographic factors cannot be ignored including limited availability of compatible males, skewed sex ratios, spatial structure, or even stochastic pairings by chance. Moreover, aggregation at the SRMR promotes social associations between bull sharks, ranging from stable companionships to casual acquaintances [37], which may increase the probability of re-encounters. Emerging biologging tools – including acoustic, satellite, and proximity tagging – are beginning to reveal such social dimensions, documenting group cohesion, dominance hierarchies, and coordinated movements in sharks [65, 66]. These tools offer promising avenues to test whether social preferences drive repeated pairings. Overall, our findings add a new perspective to shark reproductive ecology by suggesting that bull sharks in Fiji may combine polygamy with repeated pairings across years. While this pattern cannot be confirmed without direct behavioural observations, it points to the possibility that both genetic and social factors influence reproductive strategies in long-lived predators, with implications for population genetics, reproductive success, and conservation within Fiji’s diverse seascapes.

The low effective population size, together with reduced genetic diversity and evidence of kinship, these findings point to a vulnerable population status for bull sharks in Fiji. Moreover, previous genomic analyses had already revealed clear partitioning between the Fiji population and other locations at both regional and global scales, along with lower expected heterozygosity, consistent with demographic isolation [23, 35]. Combined, these results suggest that bull sharks in Fiji may face an elevated local extinction risk. Effective conservation will require large, well-enforced marine protected areas (MPAs) that safeguard all life stages and critical habitats [67]. Genetically isolated populations, such as those in Fiji, may also benefit from targeted protection of large, fecund individuals [68]. Our results further reveal fine-scale connectivity, showing that riverine habitats and the SRMR are genetically linked. The SRMR, Fiji’s first national marine park and a designated no-take zone, provides a foundation, but future MPAs should extend to coastal corridors along the south coast and protect rivers and estuaries that serve as shark parturition sites, aligning management with species’ life history and spatial ecology.

Our study meets the best-practice guidelines for kinship and relatedness research in elasmobranchs as outlined by Amini, Feutry [5], with 296 bull sharks genotyped at thousands of SNPs and sampled over a decade across multiple rivers and cohorts. Future research should prioritize systematic sampling of bull sharks from additional sites in Fiji (e.g., Vanua Levu, Yasawa group), reduced biases in adult sampling to include both sexes and bold individuals [69] and investigate the potential role of natal philopatry in this species over the long term. The evidence for multi-year pairings further suggests that reproductive behaviours in bull sharks may be more structured and persistent than previously assumed.

## Acknowledgments

We thank Gauthier Mescam, Franziska Genter, and Pascal Fluekiger, as well as the volunteers from Projects Abroad, for their assistance with field work and sample collection. We are grateful to Saki Savaraqa and Sekove Raikabu, traditional landowners in the Rewa district, for their knowledge and support during the bull shark tagging surveys. We further acknowledge the staff of Beqa Adventure Divers in Fiji, and Mike Neumann, for his continued advice during the study.

## Ethics Statement

Handling procedures of live shark specimens were approved under the “Animal Ethics Committee” section of the University of the South Pacific Research Committee and performed in accordance with relevant guidelines and regulations. All shipping procedures of bull shark tissue samples were conducted under the relevant import and export permits issued by the Australian Government, Department of Agriculture and Water Resources to Diversity Arrays Technology Pty Ltd, Canberra, Australia and the Ministry of Fisheries and Forests in Fiji. Furthermore, handling of live bull sharks at the SRMR were approved under the Research Permit issued by the Fiji Ministry of Education, Heritage and Arts to Beqa Adventure Divers and performed in accordance with relevant guidelines and regulations. During the study period, KG hold a student permit issued by the Fiji Immigration Department. Verbal consent was obtained from the Fiji Department of Fisheries in Nausori, Wainibokasi, Walu Bay, and from provincial councils before the study started and offices were informed about updates on a regular basis.

## Data Accessibility

The datasets supporting this article have been uploaded to figshare.com and can be accessed via this private link: https://figshare.com/s/bf575ea5856d3a8e66e4

## Author Contributions

KG, SA, JB, and CR designed the study. CR wrote the project proposal, and CR and KG obtained funding. KG collected bull shark samples from the Rewa River and coordinated the collection of samples from the Navua and Sigatoka Rivers as part of her PhD research at the University of the South Pacific, Suva, Fiji. JB provided samples of adult bull sharks. LB performed the kinship analyses and LB and KG created the figures. FDD validated the results. KG, LB, and CR wrote the first draft of the manuscript. All authors contributed to the final version and approved the submitted manuscript.

## Supplementary Materials

**Table S1.**
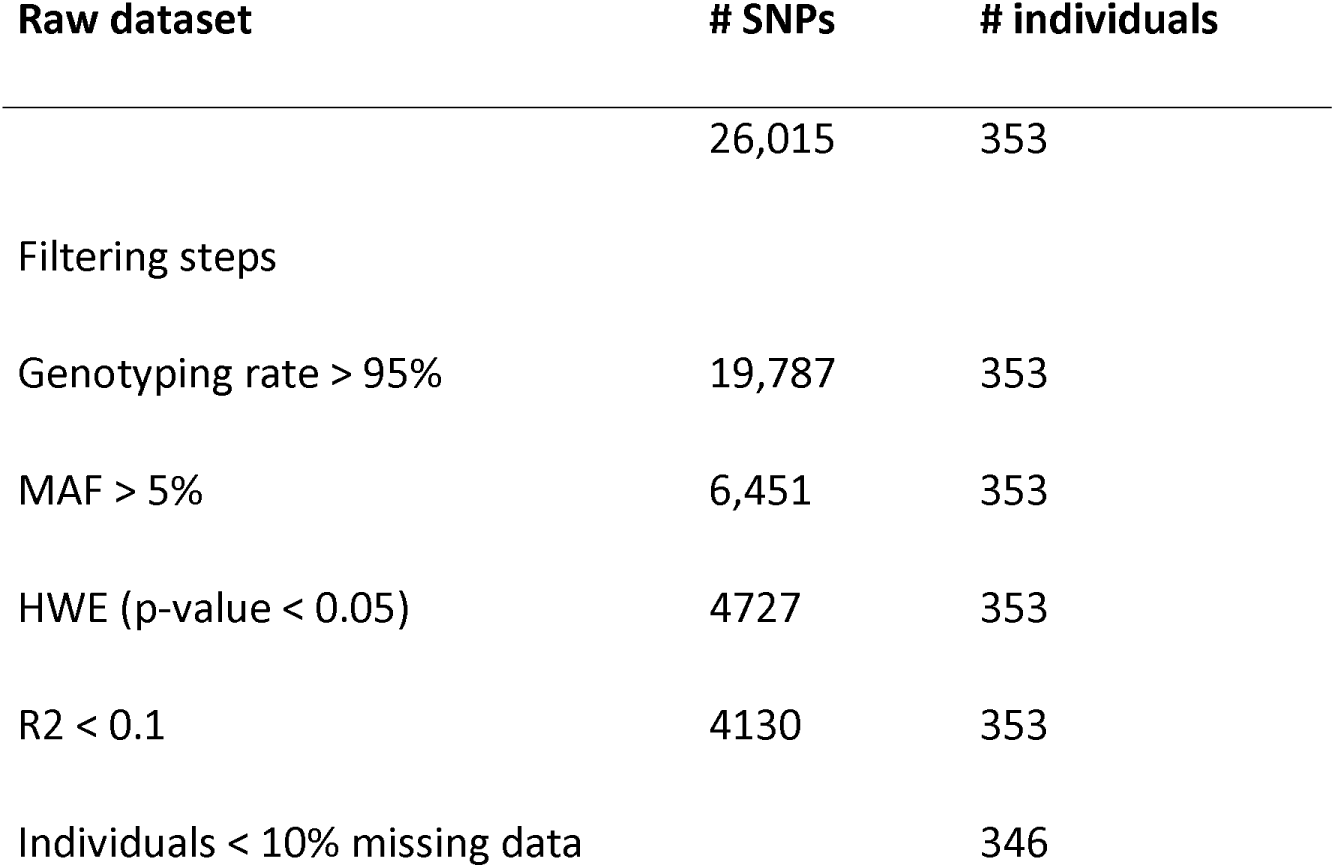
Filtering steps.

| **Raw dataset** | **# SNPs** | **# individuals** |
| --- | --- | --- |
|  | 26,015 | 353 |
| Filtering steps |  |  |
| Genotyping rate > 95% | 19,787 | 353 |
| MAF > 5% | 6,451 | 353 |
| HWE (p-value < 0.05) | 4727 | 353 |
| R2 < 0.1 | 4130 | 353 |
| Individuals < 10% missing data |  | 346 |

**Figure S1.**
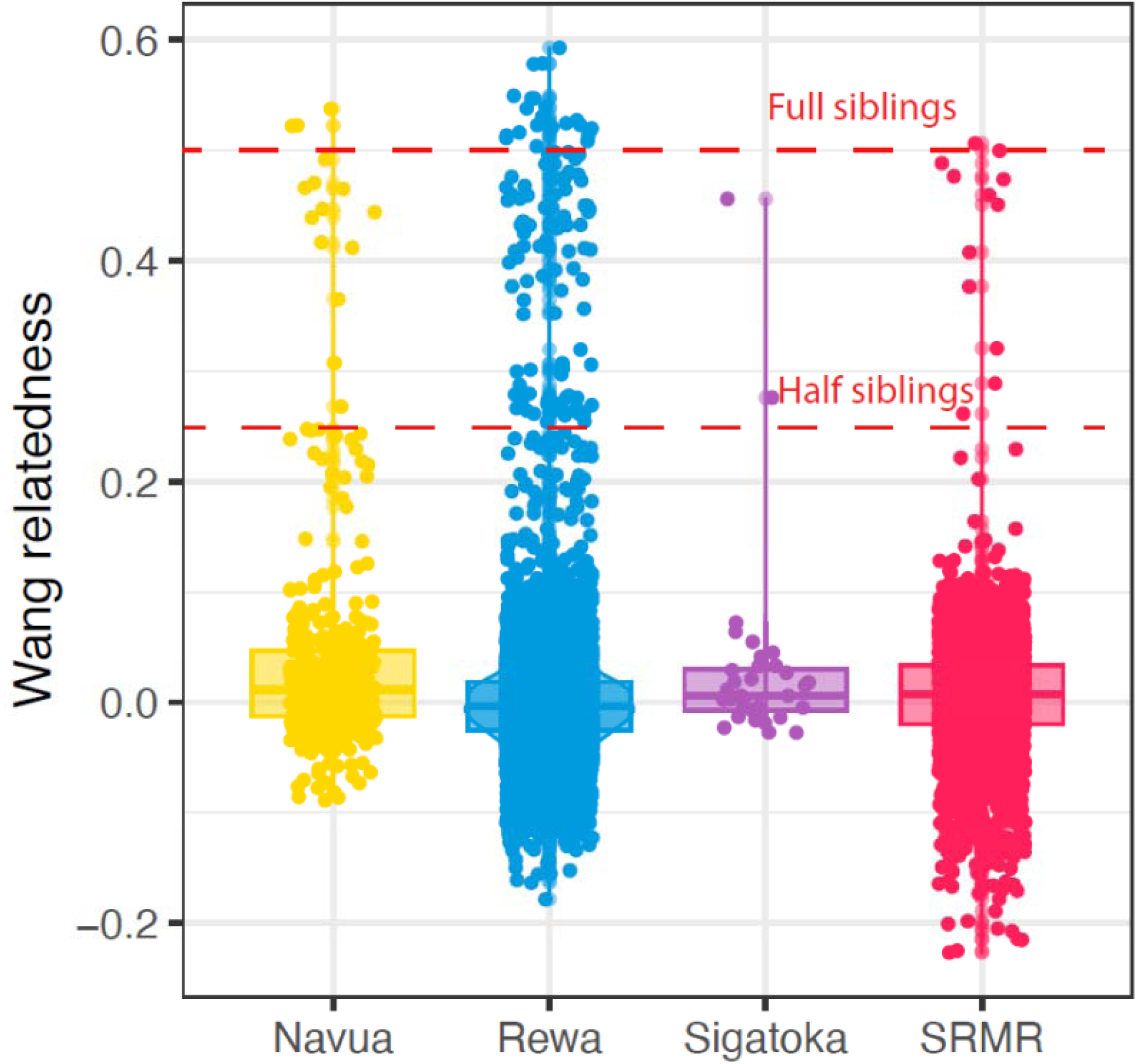
Wang estimator on pairwise genetic relatedness.

**Table S2.**
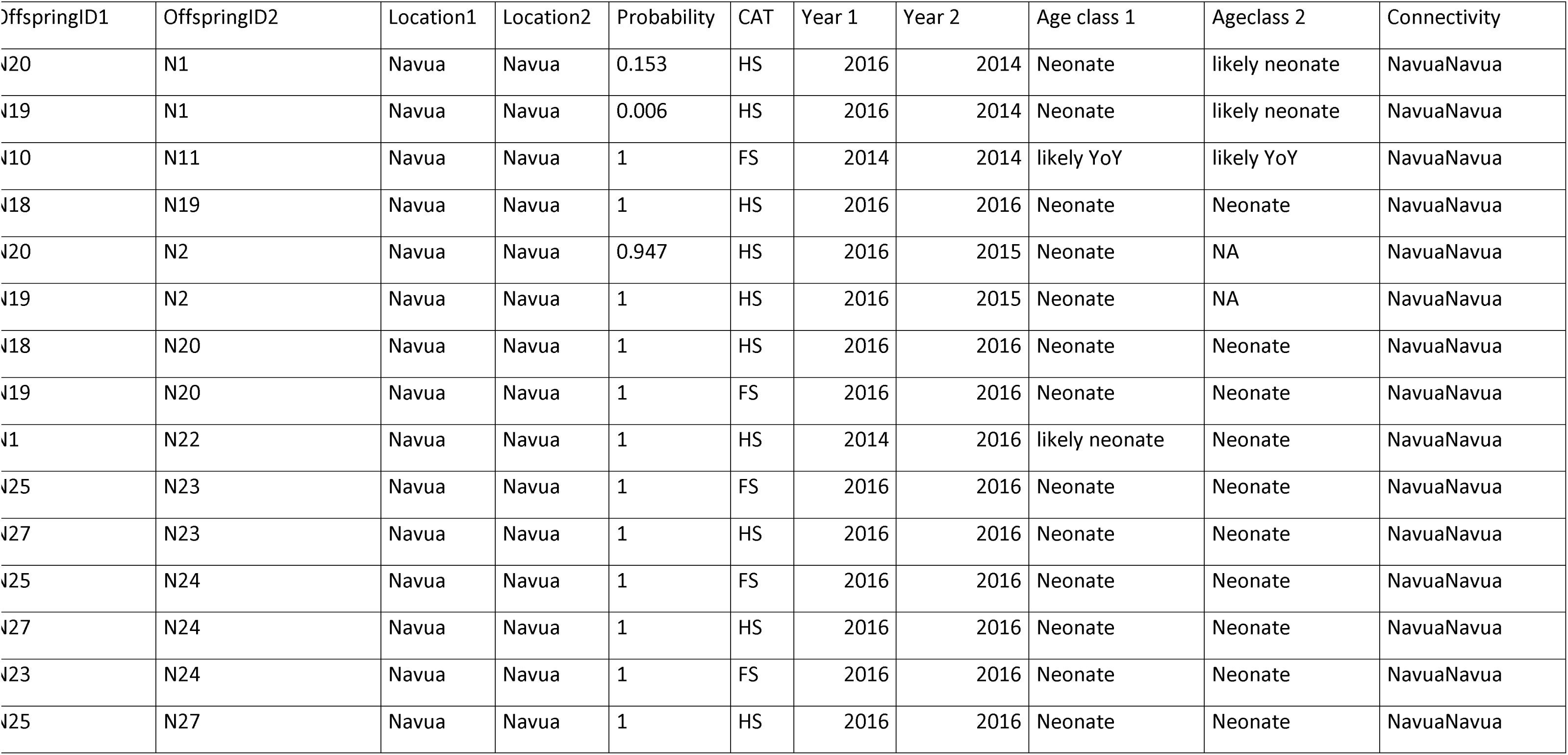
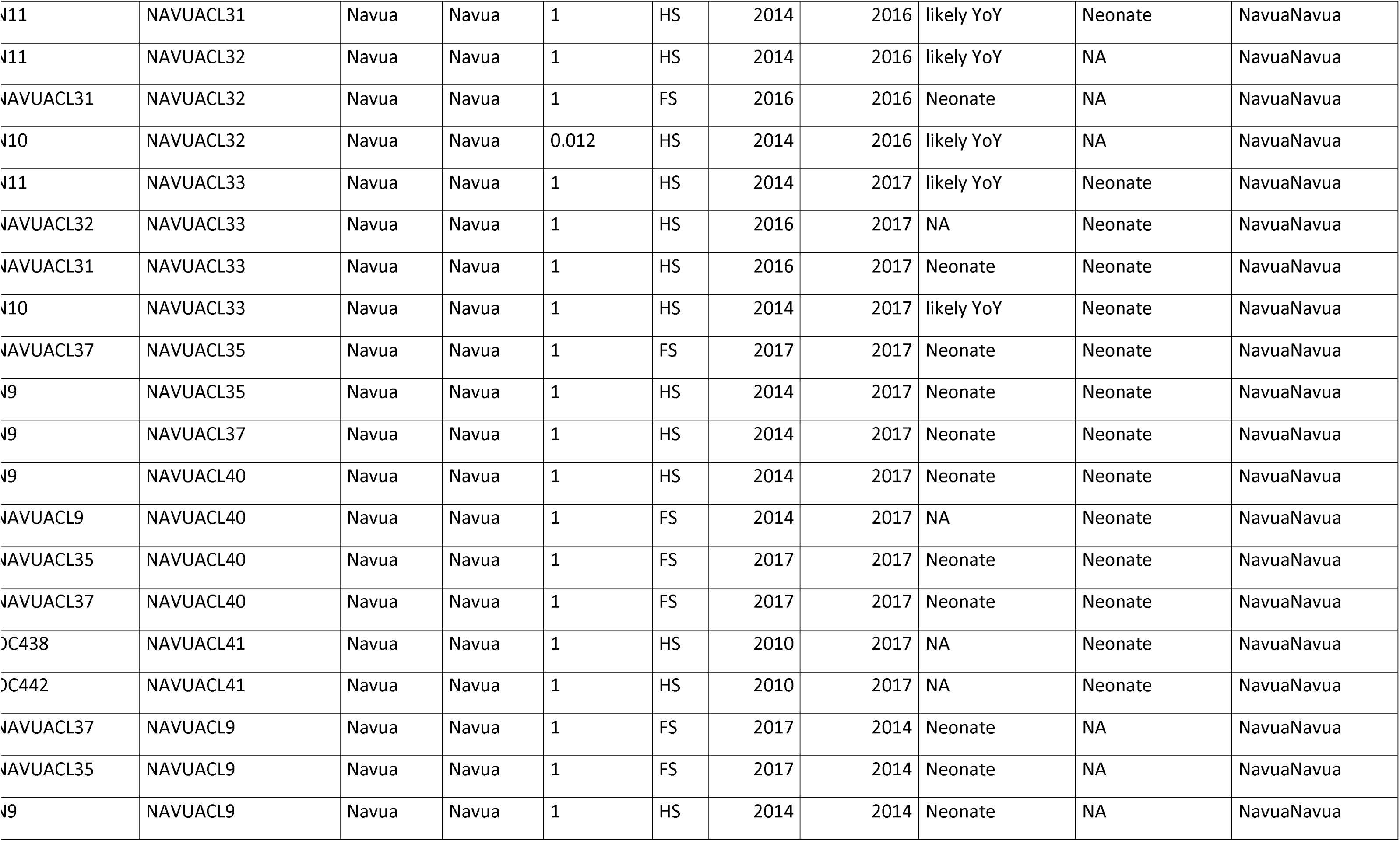
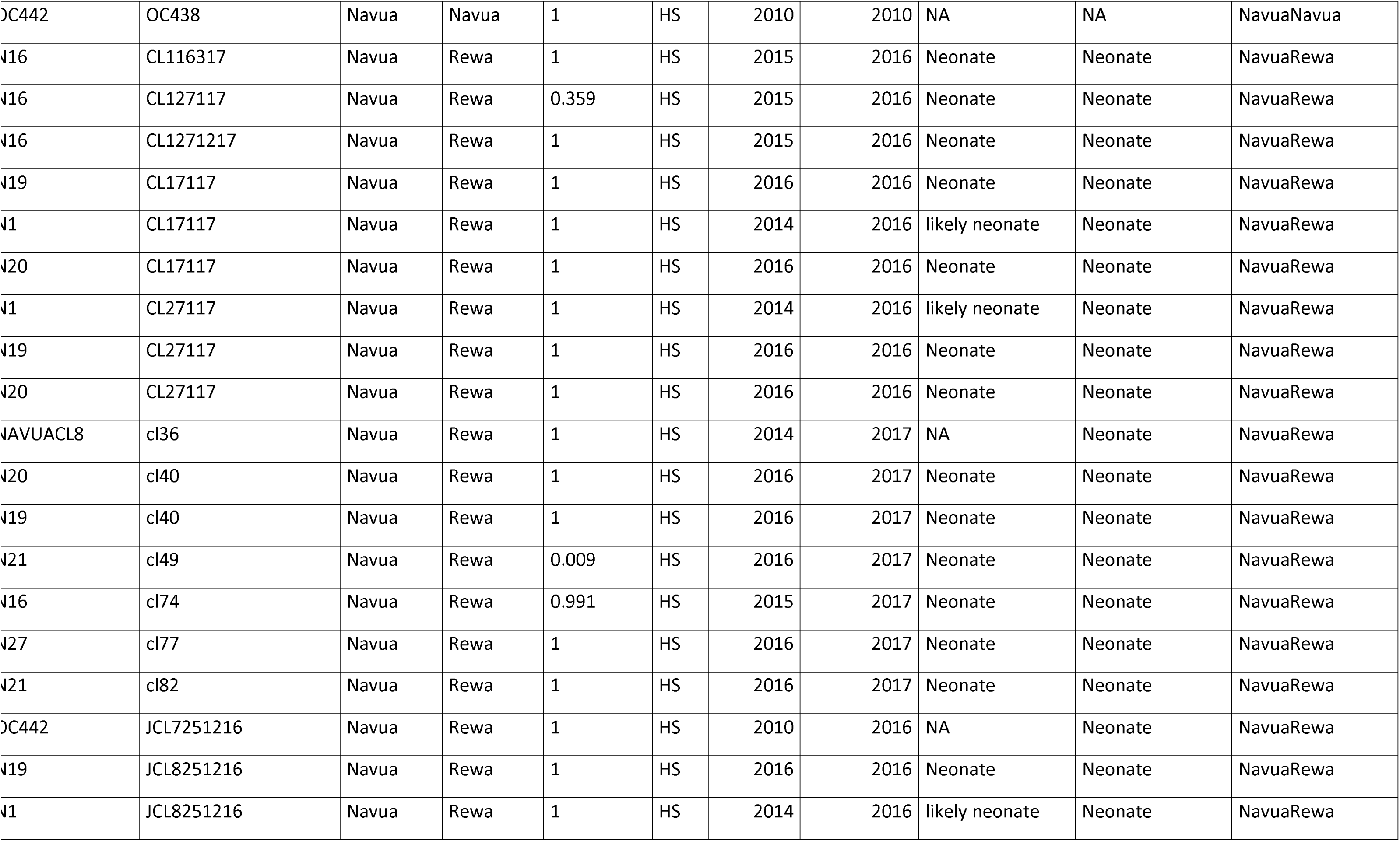
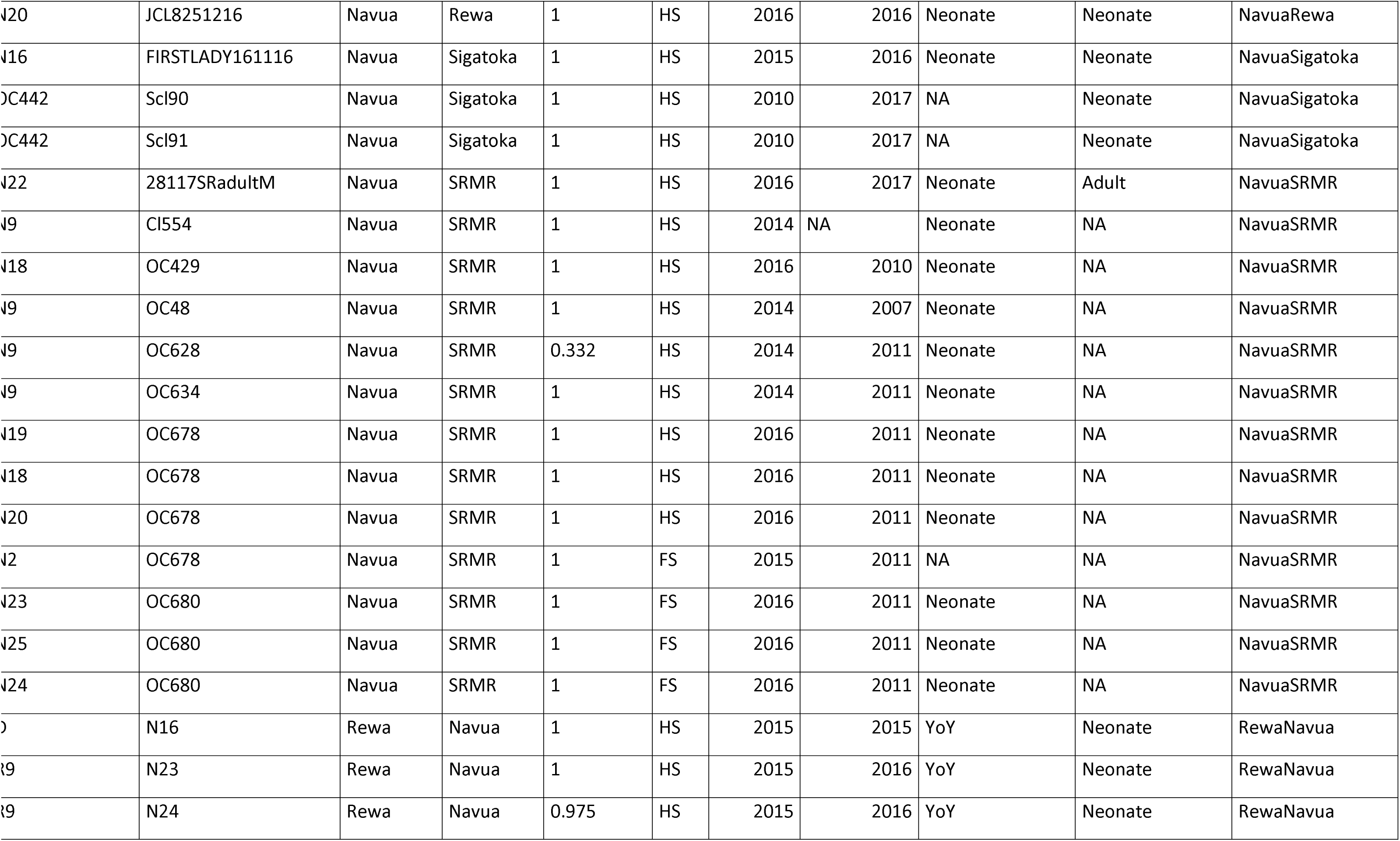
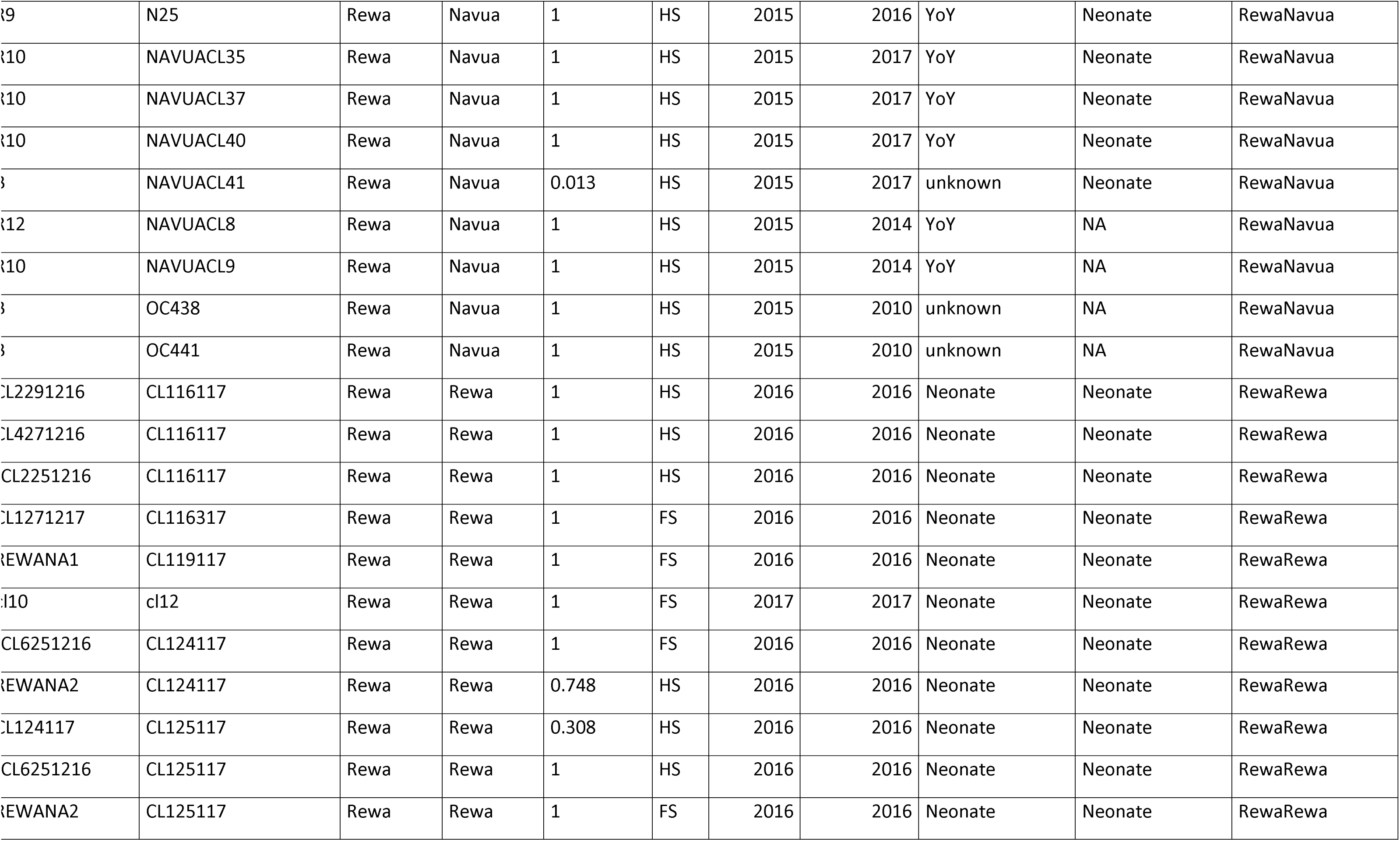
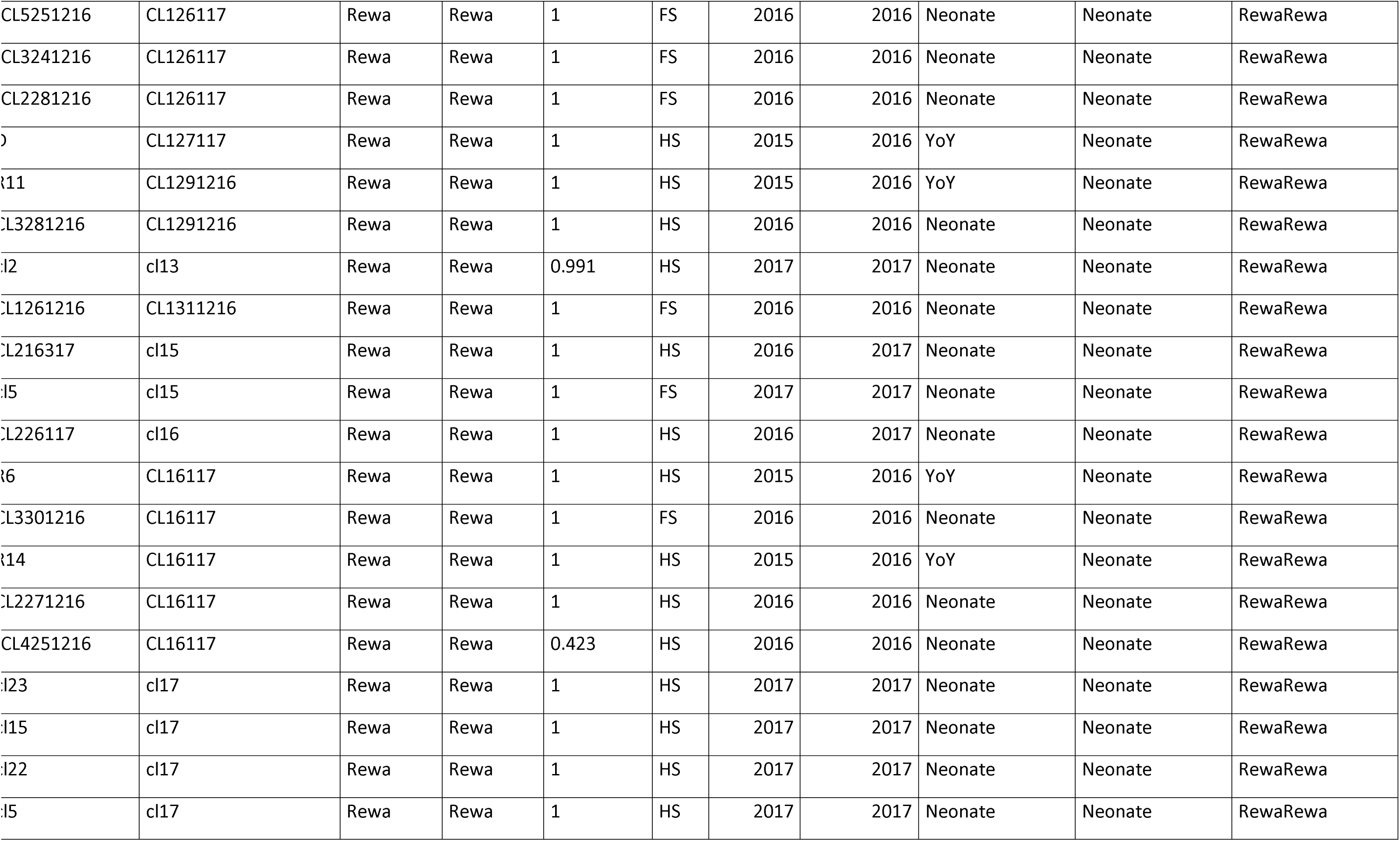
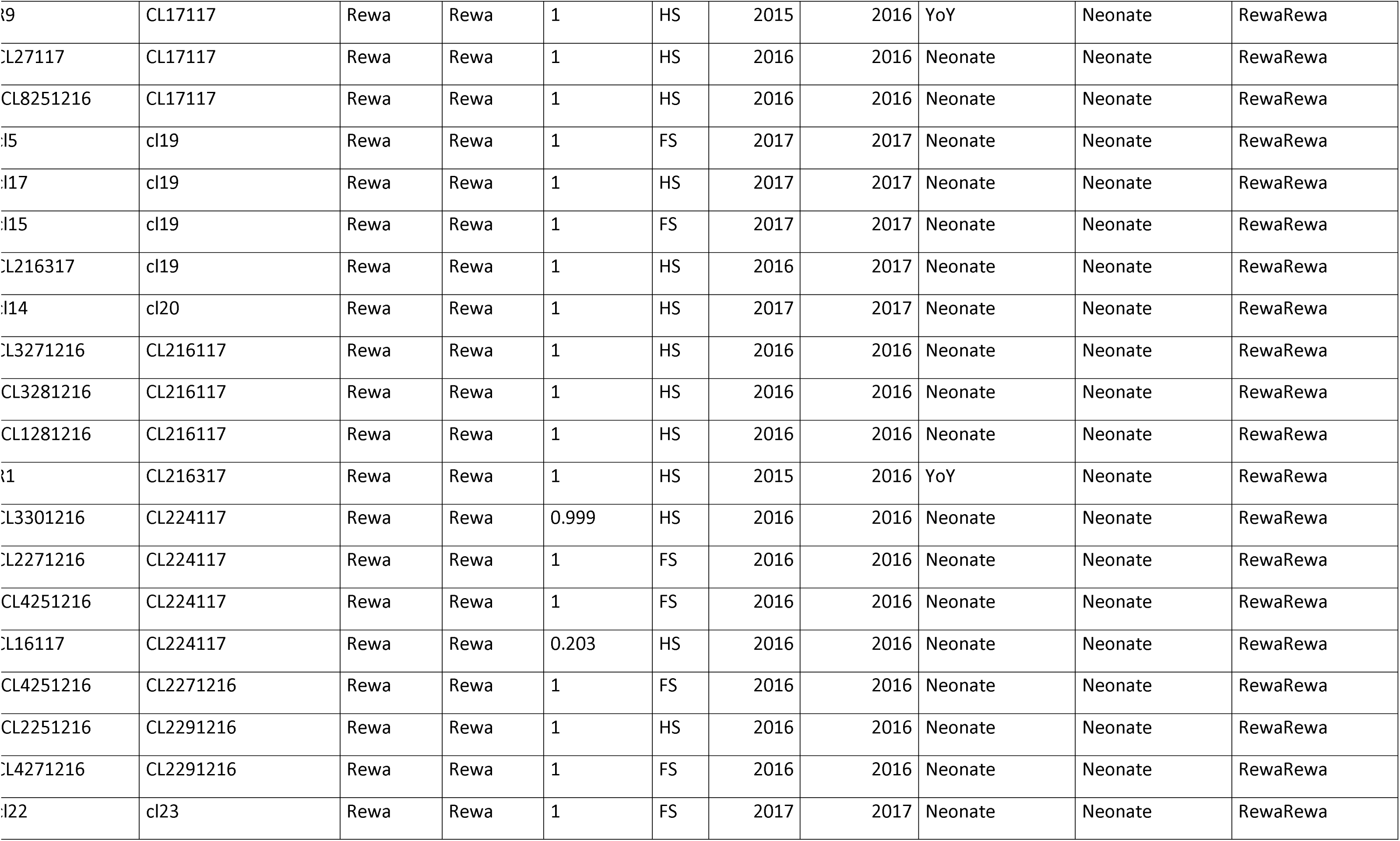
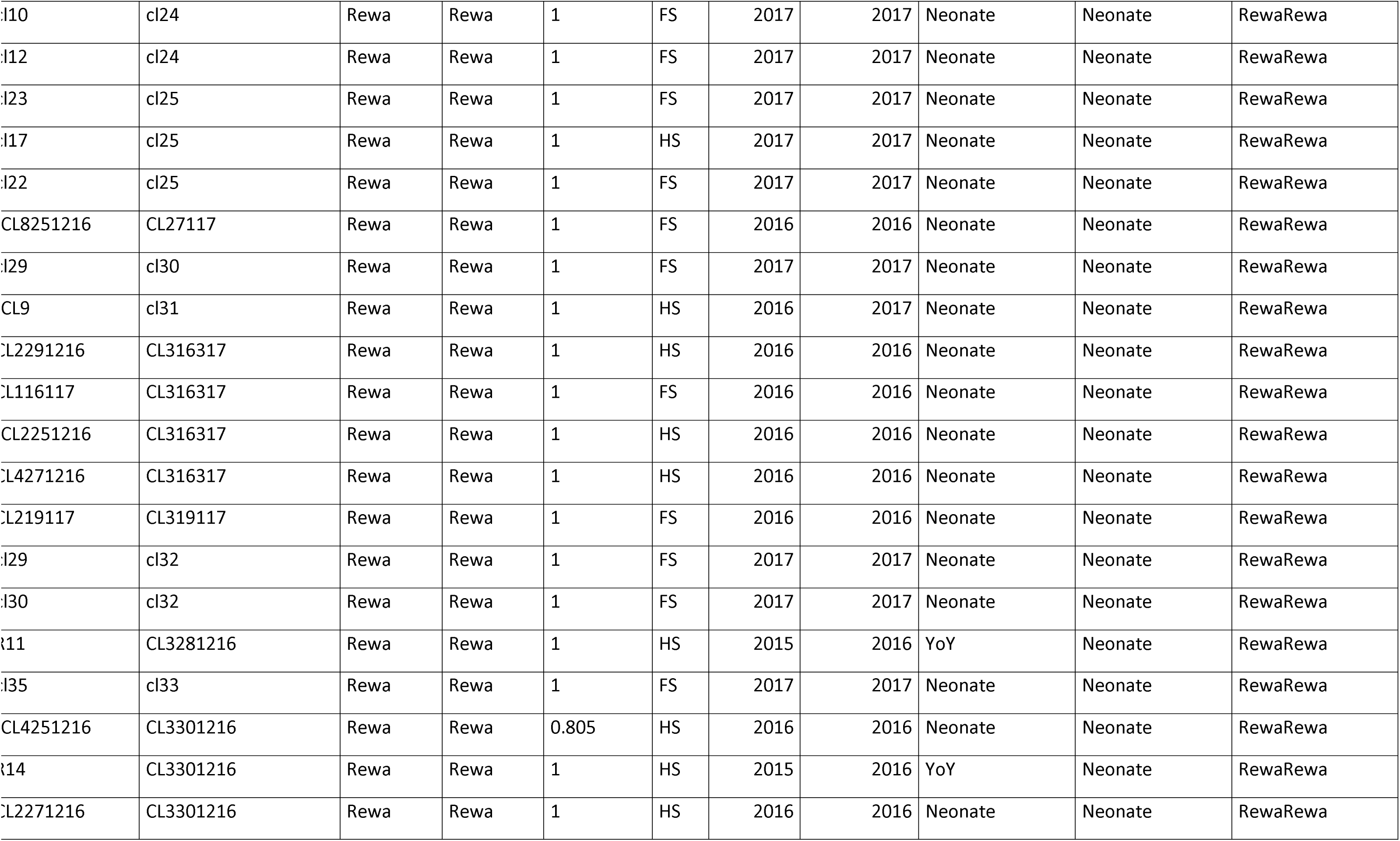
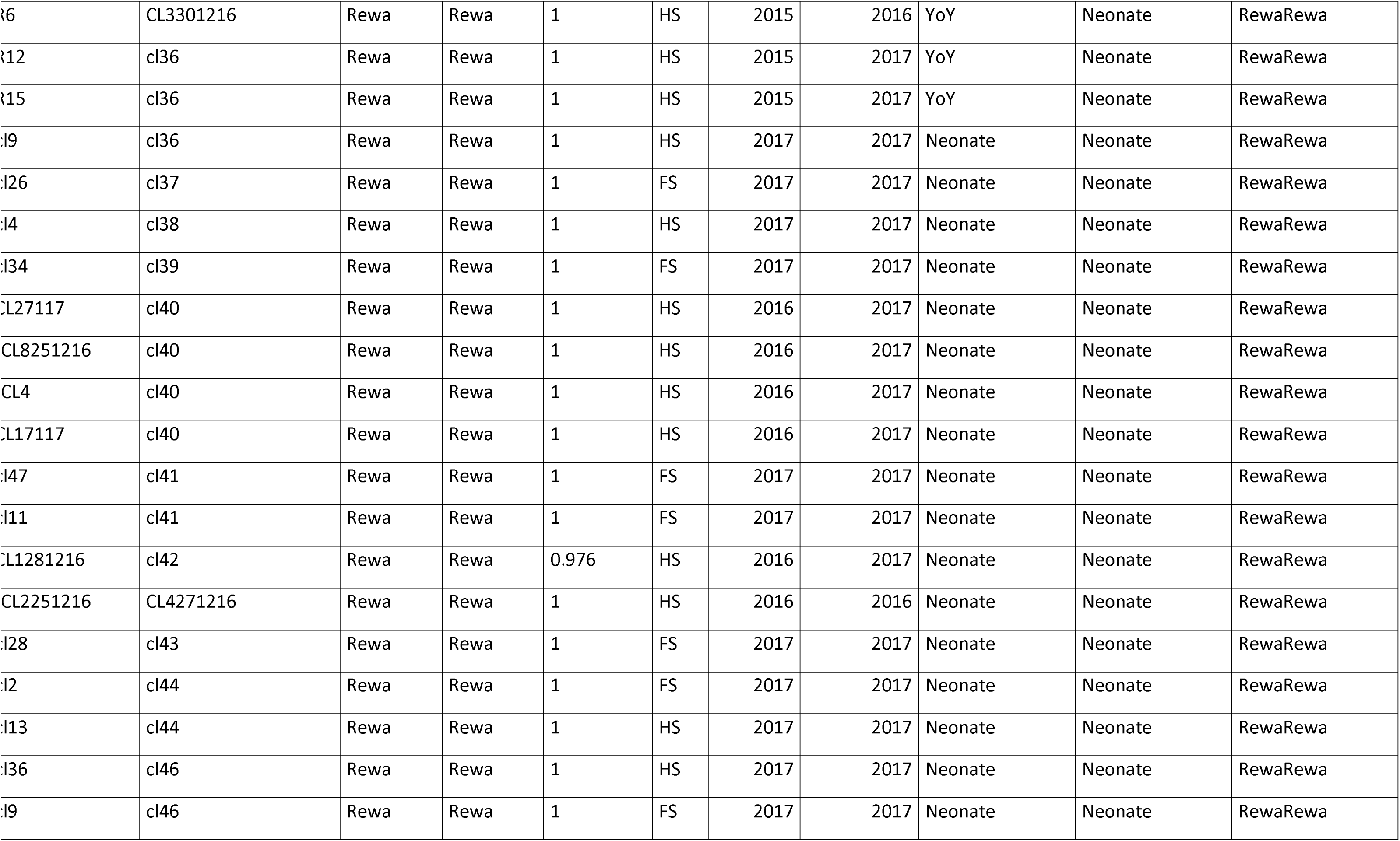
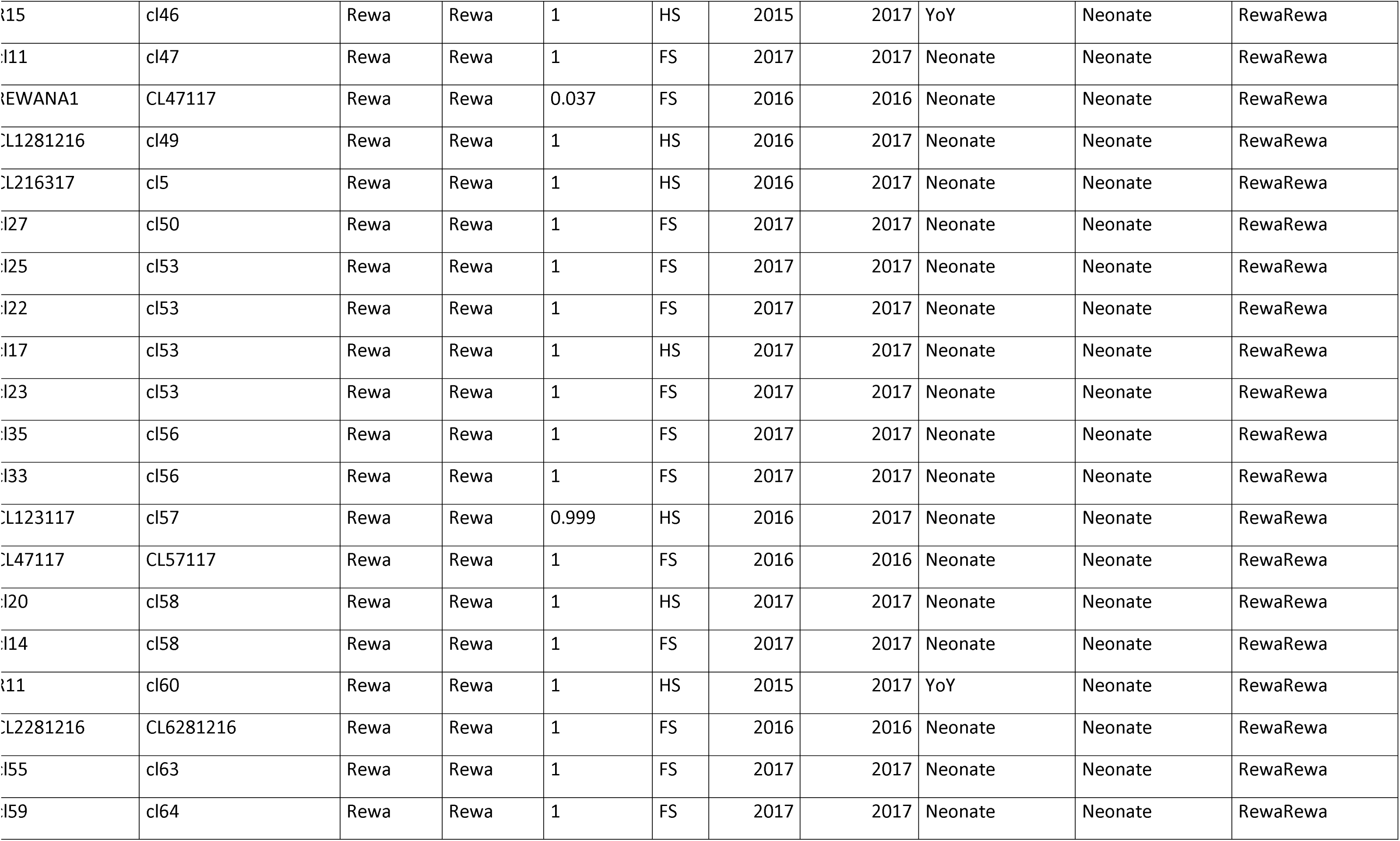
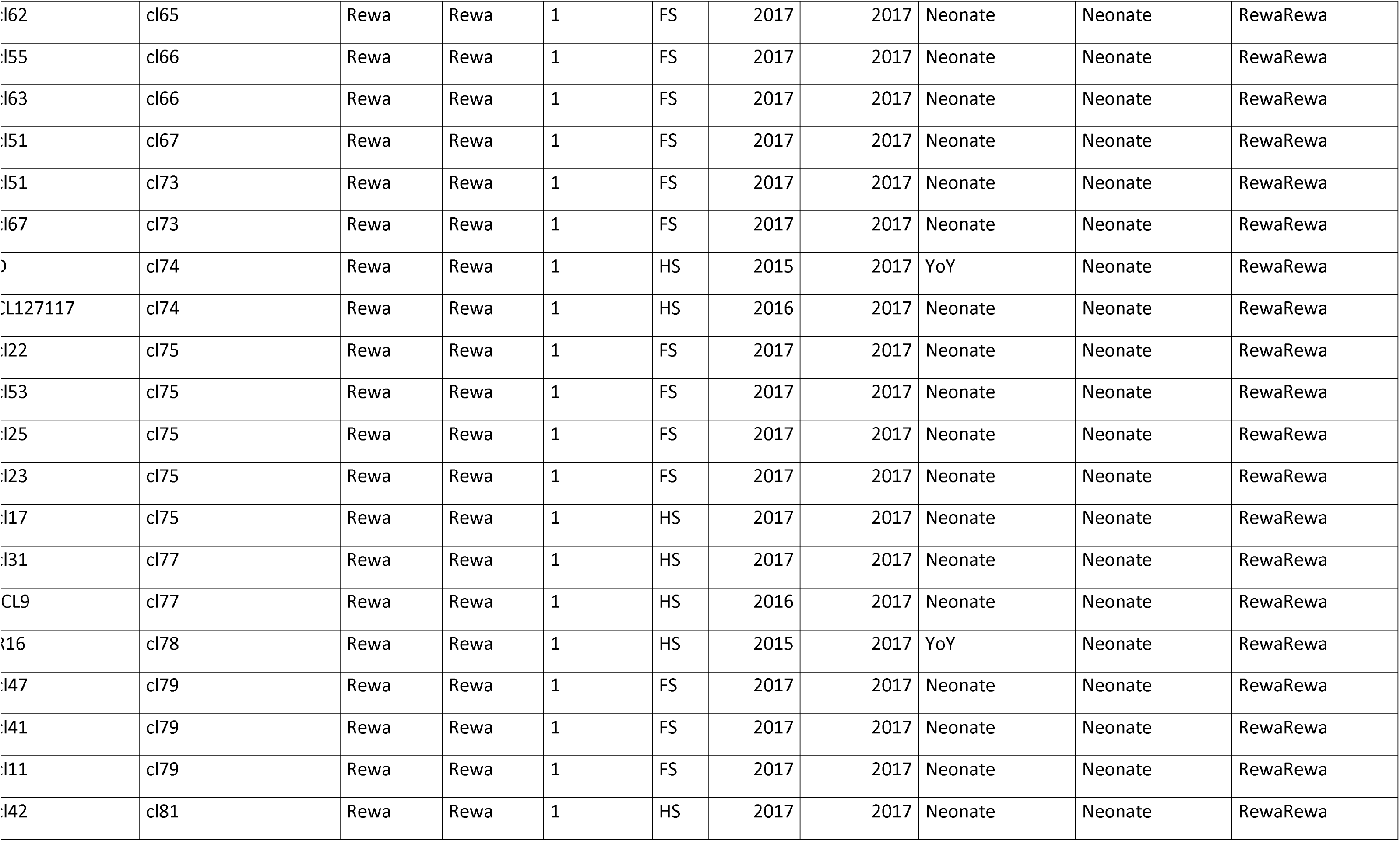
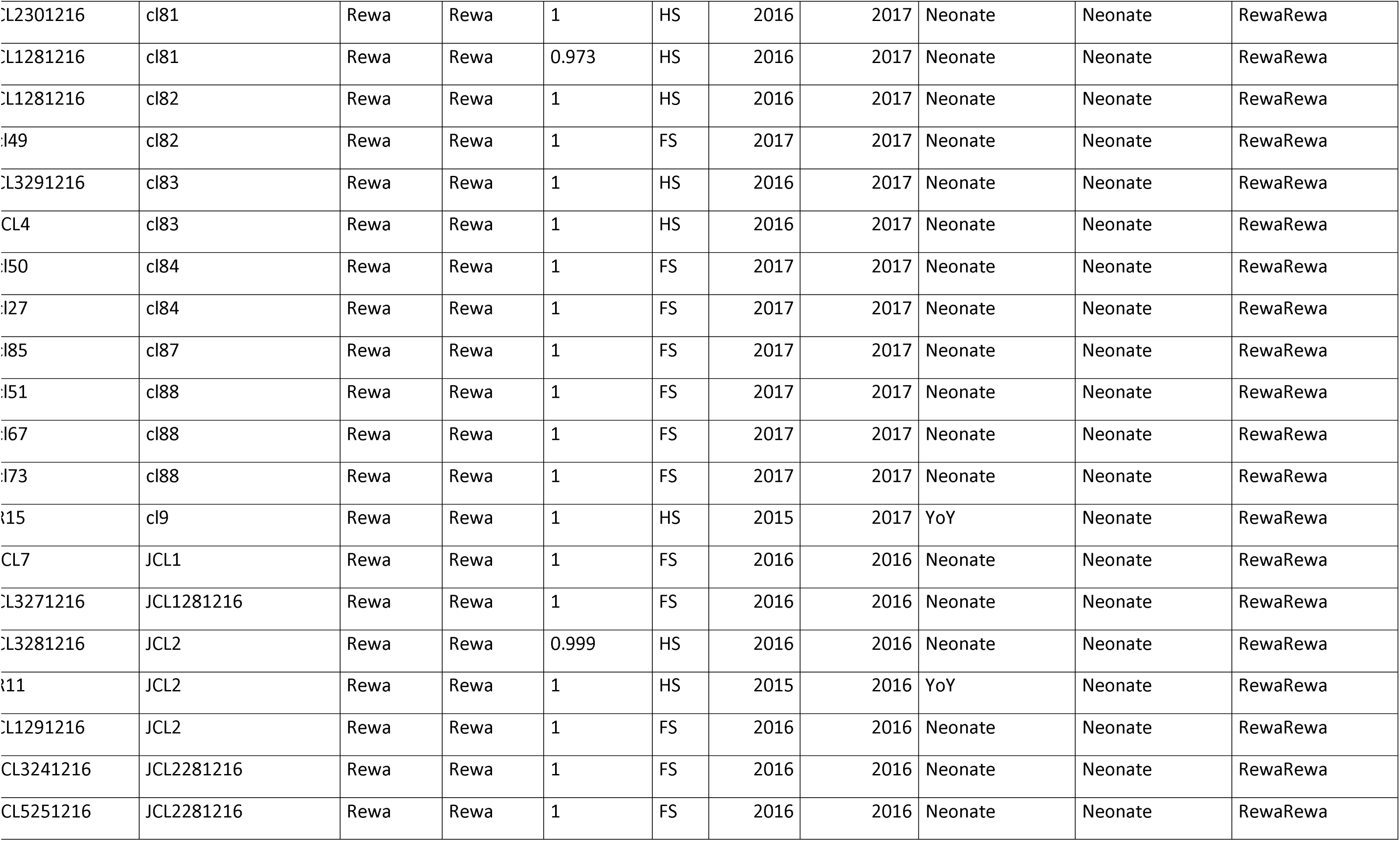
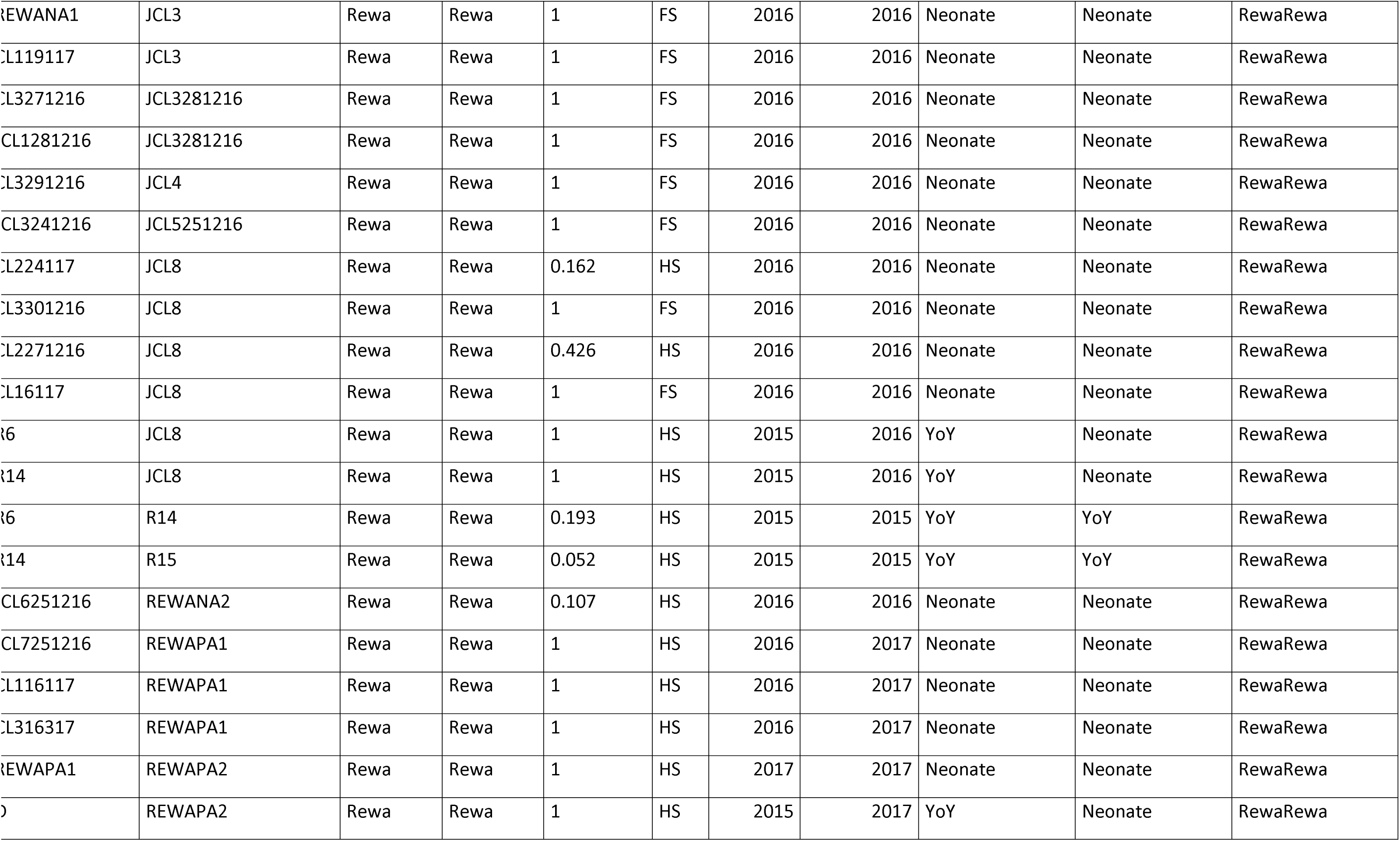
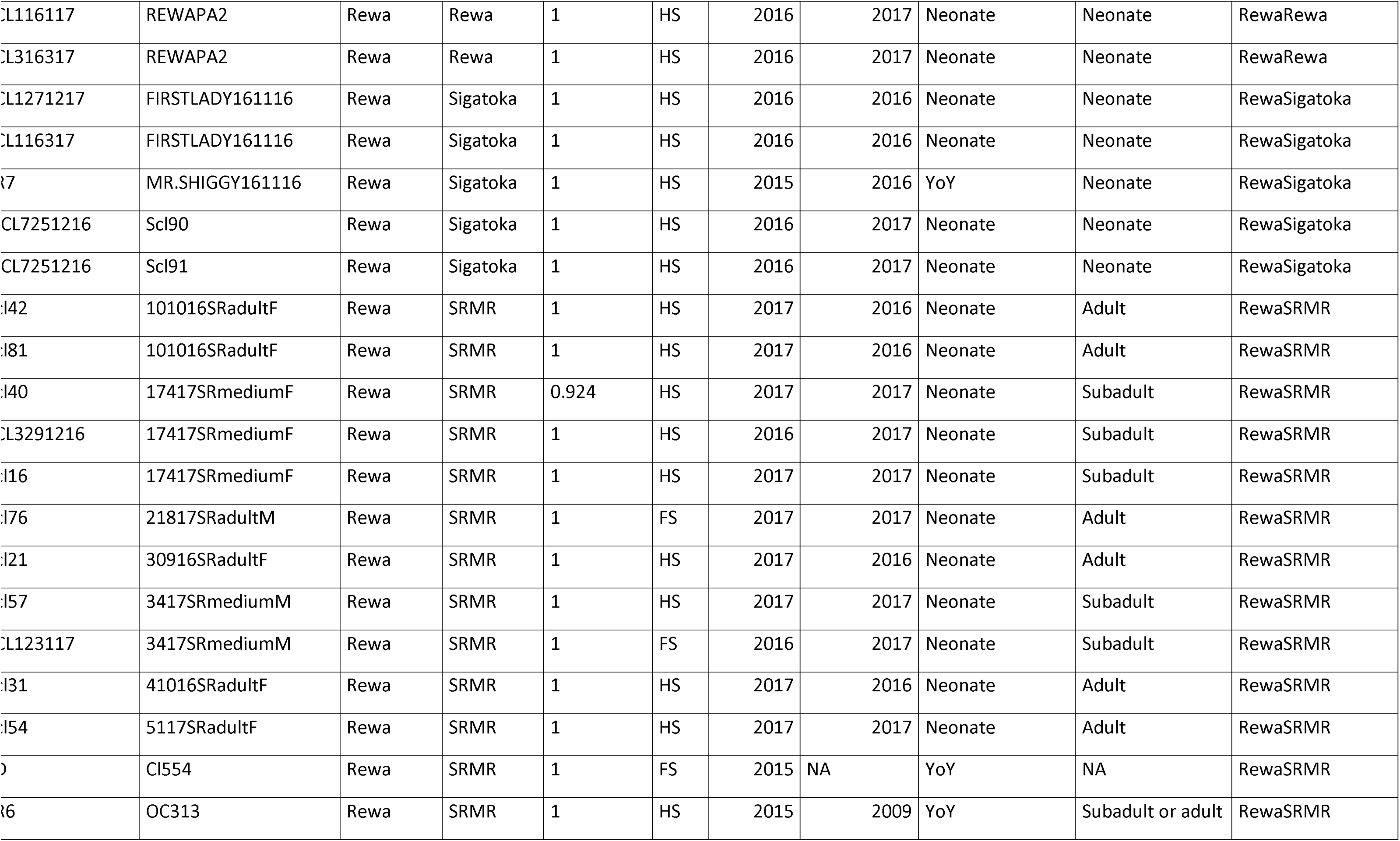
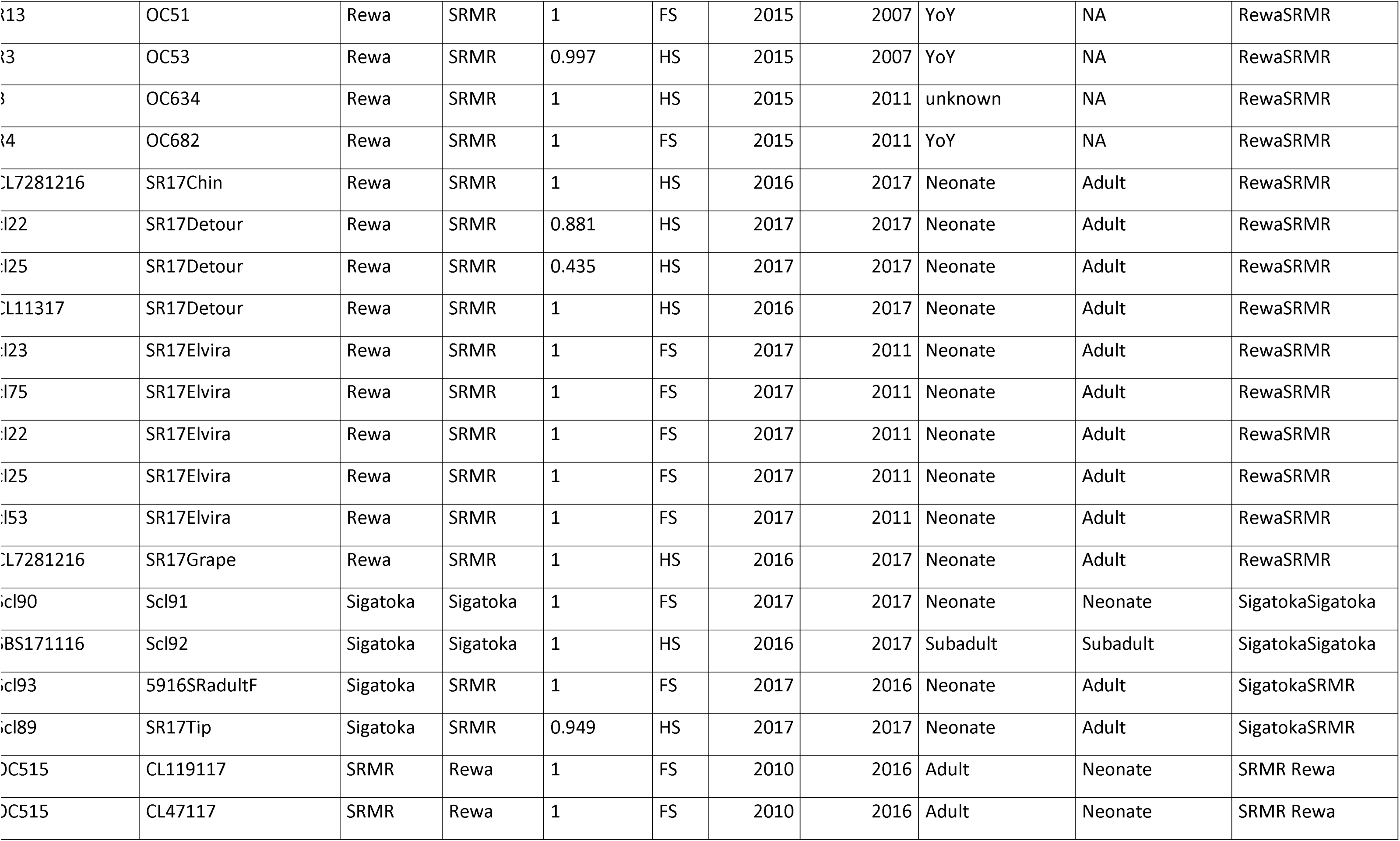
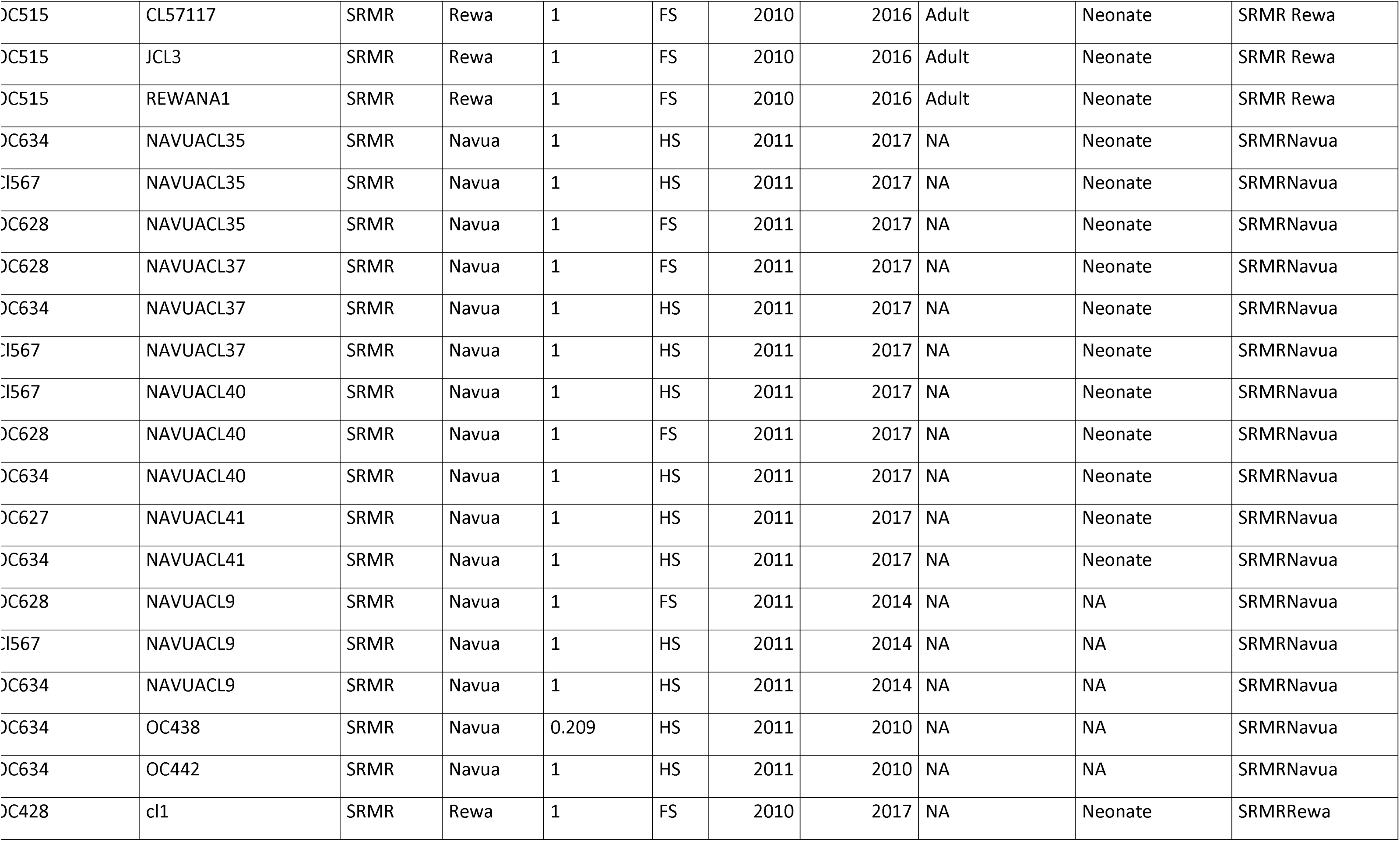
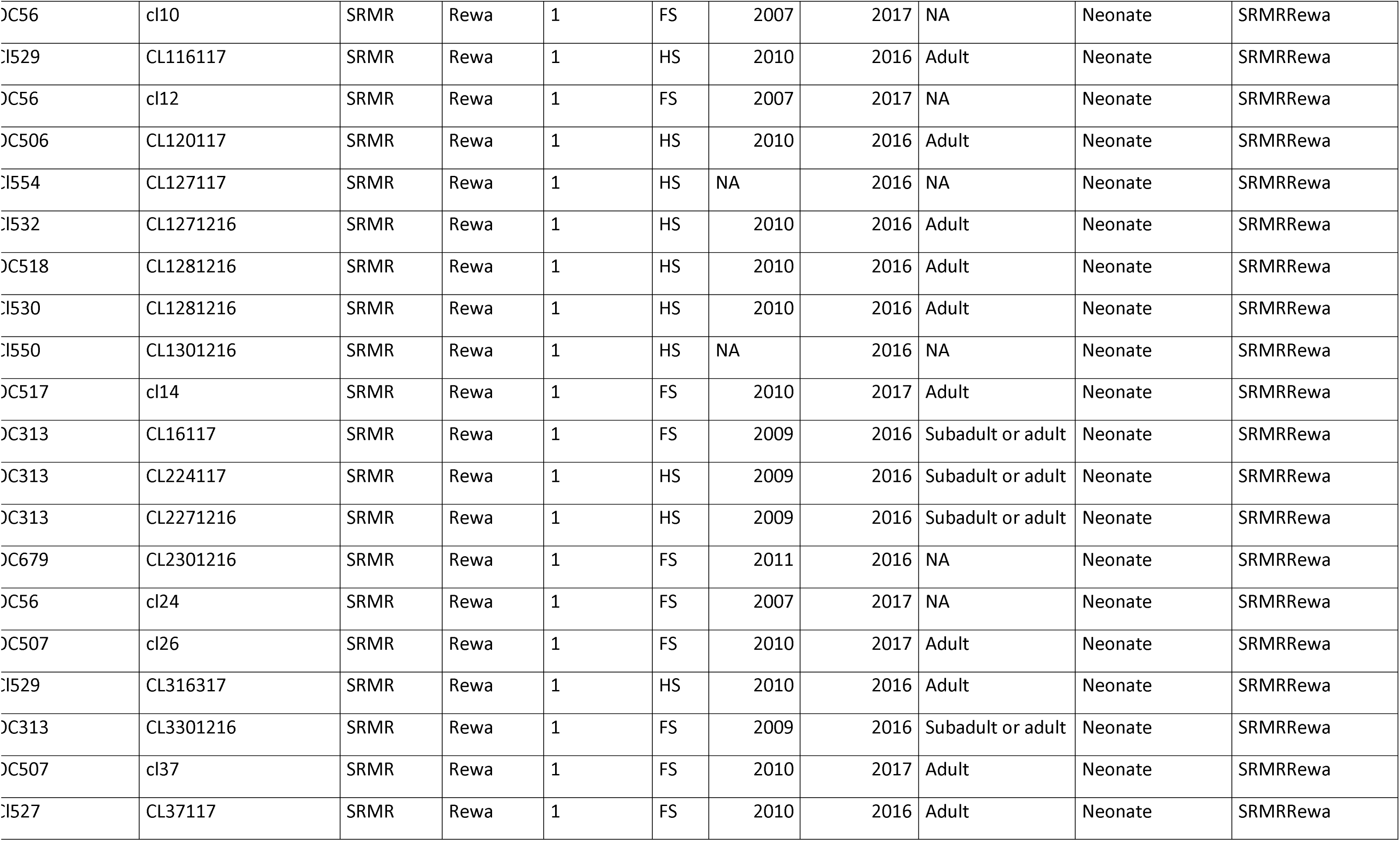
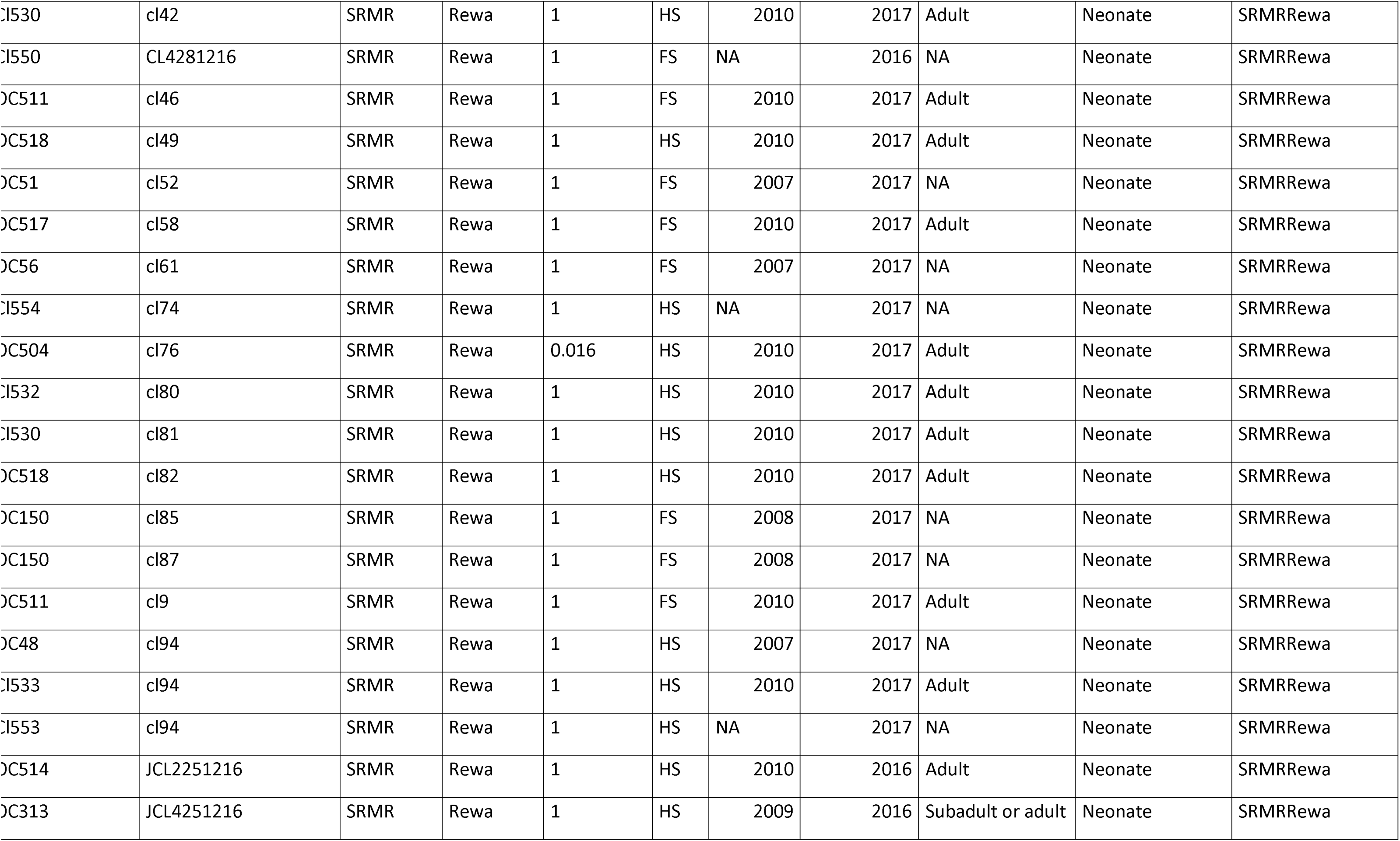
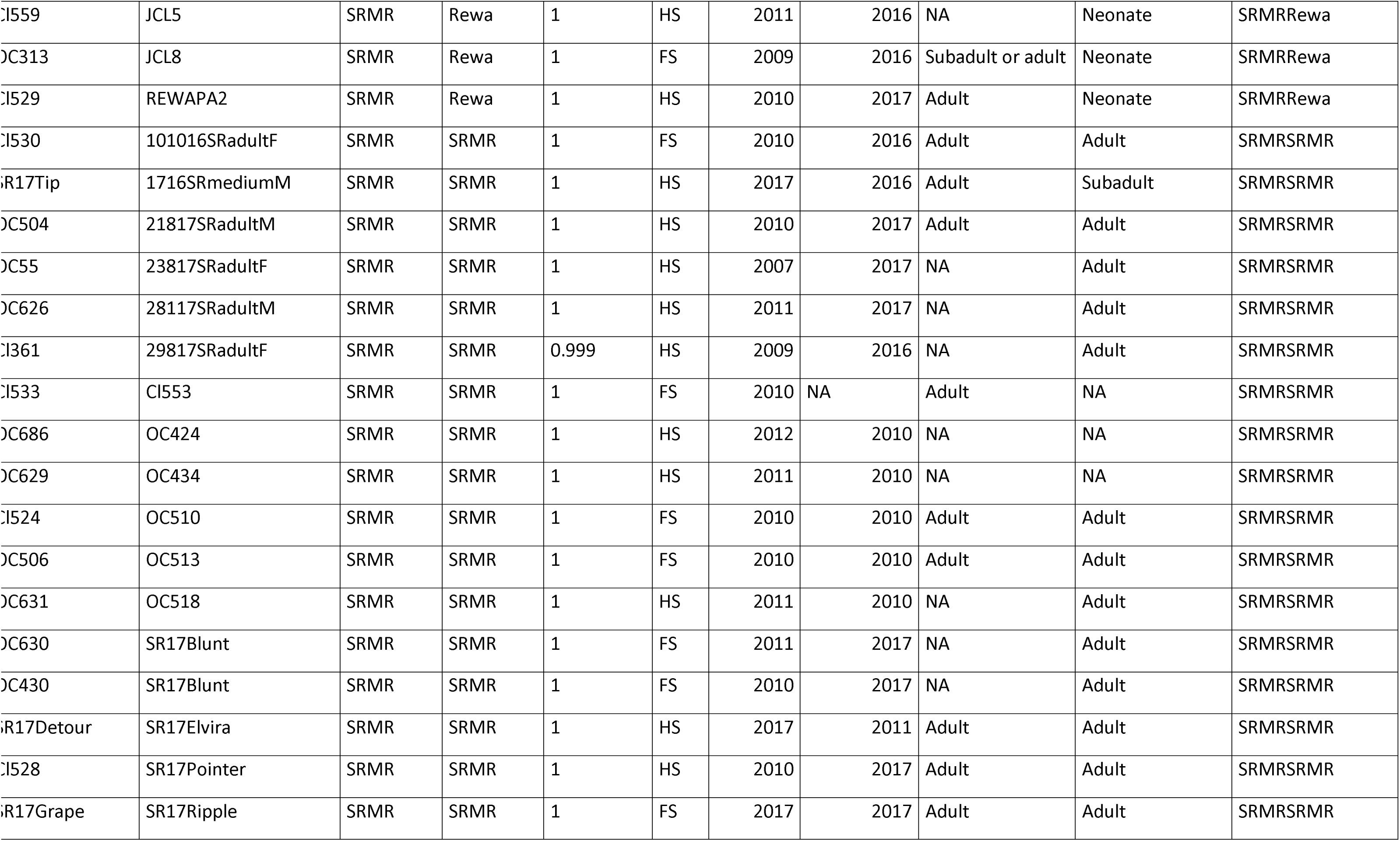
All family clusters 2.

## References

[1] Simpfendorfer, C.A., Heithaus, M.R., Heupel, M.R., MacNeil, M.A., Meekan, M., Harvey, E., Sherman, C.S., Currey-Randall, L.M., Goetze, J.S. & Kiszka, J.J. 2023 Widespread diversity deficits of coral reef sharks and rays. Science 380, 1155–1160.

[2] Cortés, E. 2000 Life history patterns and correlations in sharks. Reviews in Fisheries Science 8, 299–344.

[3] Parsons, G.R., Hoffmayer, E.R., Hendon, J.M., Bet-Sayad, W.V., Rocha, M., Arukwe, A. & Kapoor, B. 2008 A review of shark reproductive ecology: life history and evolutionary implications. Fish reproduction 1, 435–469.

[4] Speed, C.W., Field, I.C., Meekan, M.G. & Bradshaw, C.J. 2010 Complexities of coastal shark movements and their implications for management. Marine ecology progress series 408, 275–293.

[5] Amini, S.R., Feutry, P., Phillips, N.M. & Kyne, P.M. 2025 A review of the utility and application of relatedness and kinship in elasmobranchs. Reviews in Fish Biology and Fisheries. (doi:10.1007/s11160-025-09983-2).

[6] Städele, V. & Vigilant, L. 2016 Strategies for determining kinship in wild populations using genetic data. In Ecology and Evolution (

[7] Mourier, J. & Planes, S. 2013 Direct genetic evidence for reproductive philopatry and associated fine-scale migrations in female blacktip reef sharks (Carcharhinus melanopterus) in French Polynesia. Molecular Ecology 22. (doi:10.1111/mec.12103).

[8] Morgan, Barry, S., Gulak, S.J.B. & Drymon, J.M. 2022 Is blood thicker than water? Relatedness and kinship in Bull Sharks (Carcharhinus leucas). Florida Scientist 85, 59–59.

[9] Feldheim, K.A., Gruber, S.H., Dibattista, J.D., Babcock, E.A., Kessel, S.T., Hendry, A.P., Pikitch, E.K., Ashley, M.V. & Chapman, D.D. 2014 Two decades of genetic profiling yields first evidence of natal philopatry and long-term fidelity to parturition sites in sharks. Molecular Ecology 23. (doi:10.1111/mec.12583).

[10] DiBattista, J.D., Feldheim, K.A., Thibert-Plante, X., Gruber, S.H. & Hendry, A.P. 2008 A genetic assessment of polyandry and breeding-site fidelity in lemon sharks. Molecular Ecology 17. (doi:10.1111/j.1365-294X.2008.03833.x).

[11] Patterson, T.A., Hillary, R.M., Kyne, P.M., Pillans, R.D., Gunasekera, R.M., Marthick, J.R., Johnson, G.J. & Feutry, P. 2022 Rapid assessment of adult abundance and demographic connectivity from juvenile kin pairs in a critically endangered species. Science Advances 8, eadd1679.

[12] Guttridge, T.L., Gruber, S.H., Franks, B.R., Kessel, S.T., Gledhill, K.S., Uphill, J., Krause, J. & Sims, D.W. 2012 Deep danger: Intra-specific predation risk influences habitat use and aggregation formation of juvenile lemon sharks Negaprion brevirostris. Marine Ecology Progress Series 445. (doi:10.3354/meps09423).

[13] Larson, S., Christiansen, J., Griffing, D., Ashe, J., Lowry, D. & Andrews, K. 2011 Relatedness and polyandry of sixgill sharks, Hexanchus griseus, in an urban estuary. Conservation Genetics 12. (doi:10.1007/s10592-010-0174-9).

[14] Hueter, R.E., Heupel, M., Heist, E. & Keeney, D. 2005 Evidence of philopatry in sharks and implications for the management of shark fisheries. Journal of northwest atlantic fishery Science 35, 239–247.

[15] Gausmann, P. 2021 Synopsis of global fresh and brackish water occurrences of the bull shark Carcharhinus leucas Valenciennes, 1839 (Pisces: Carcharhinidae), with comments on distribution and habitat use. Integrative Systematics 4. (doi:10.18476/2021.423083).

[16] Smoothey, A.F., Lee, K.A. & Peddemors, V.M. 2019 Long-term patterns of abundance, residency and movements of bull sharks (Carcharhinus leucas) in Sydney Harbour, Australia. Scientific Reports 9. (doi:10.1038/s41598-019-54365-x).

[17] Trystram, C., Rogers, K.M., Soria, M. & Jaquemet, S. 2017 Feeding patterns of two sympatric shark predators in coastal ecosystems of an oceanic island. Canadian Journal of Fisheries and Aquatic Sciences 74. (doi:10.1139/cjfas-2016-0105).

[18] TinHan, T.C. & Wells, R.J.D. 2021 Spatial and Ontogenetic Patterns in the Trophic Ecology of Juvenile Bull Sharks (Carcharhinus leucas) From the Northwest Gulf of Mexico. Frontiers in Marine Science 8. (doi:10.3389/fmars.2021.664316).

[19] Pirog, A., Magalon, H., Poirout, T. & Jaquemet, S. 2019 Reproductive biology, multiple paternity and polyandry of the bull shark Carcharhinus leucas. Journal of Fish Biology 95, 1195–1206.

[20] Werry, J., Lee, S., Otway, N., Hu, Y. & Sumpton, W. 2011 A multi-faceted approach for quantifying the estuarine–nearshore transition in the life cycle of the bull shark, Carcharhinus leucas. Marine and Freshwater Research 62, 1421–1431.

[21] Matich, P. & Heithaus, M.R. 2015 Individual variation in ontogenetic niche shifts in habitat use and movement patterns of a large estuarine predator (Carcharhinus leucas). Oecologia 178, 347–359.

[22] Heupel, M.R. & Simpfendorfer, C.A. 2011 Estuarine nursery areas provide a low-mortality environment for young bull sharks Carcharhinus leucas. Marine Ecology Progress Series 433, 237–244.

[23] Devloo-Delva, F., Burridge, C.P., Kyne, P.M., Brunnschweiler, J.M., Chapman, D.D., Charvet, P., Chen, X., Cliff, G., Daly, R., Drymon, J.M., et al. 2023 From rivers to ocean basins: The role of ocean barriers and philopatry in the genetic structuring of a cosmopolitan coastal predator. Ecology and Evolution 13. (doi:10.1002/ece3.9837).

[24] Postaire, B.D., Devloo-Delva, F., Brunnschweiler, J.M., Charvet, P., Chen, X., Cliff, G., Daly, R., Drymon, J.M., Espinoza, M., Fernando, D., et al. 2024 Global genetic diversity and historical demography of the Bull Shark. Journal of Biogeography 51. (doi:10.1111/jbi.14774).

[25] Tillett, B., Meekan, M., Field, I., Thorburn, D. & Ovenden, J. 2012 Evidence for reproductive philopatry in the bull shark Carcharhinus leucas. Journal of fish biology 80, 2140–2158.

[26] Lea, J.S.E., Humphries, N.E., Clarke, C.R. & Sims, D.W. 2015 To Madagascar and back: Long-distance, return migration across open ocean by a pregnant female bull shark Carcharhinus leucas. Journal of Fish Biology 87. (doi:10.1111/jfb.12805).

[27] Sherman, C.S., Simpfendorfer, C.A., Pacoureau, N., Matsushiba, J.H., Yan, H.F., Walls, R.H., Rigby, C.L., VanderWright, W.J., Jabado, R.W. & Pollom, R.A. 2023 Half a century of rising extinction risk of coral reef sharks and rays. Nature Communications 14, 15.

[28] Matich, P., Plumlee, J.D., Bubley, W., Curtis, T.H., Drymon, J.M., Mullins, L.L., Shipley, O.N., TinHan, T.C. & Fisher, M.R. 2024 Long term effects of climate change on juvenile bull shark migratory patterns. Journal of Animal Ecology.

[29] Glaus, K.B., Adrian-Kalchhauser, I., Piovano, S., Appleyard, S.A., Brunnschweiler, J.M. & Rico, C. 2019 Fishing for profit or food? Socio-economic drivers and fishers’ attitudes towards sharks in Fiji. Marine Policy 100, 249–257.

[30] Brunnschweiler, J.M. & Marosi, N.D. 2019 Two years of impairment: Plastic packing strap on a bull shark (Carcharhinus leucas) in Fiji. Pacific Conservation Biology 26, 208–209.

[31] Rigby & Espinoza. 2020 Carcharhinus leucas: Rigby, C.L., Espinoza, M., Derrick, D., Pacoureau, N. & Dicken, M. In IUCN Red List of Threatened Species (

[32] Brunnschweiler, J.M. & Baensch, H. 2011 Seasonal and long-term changes in relative abundance of bull sharks from a tourist shark feeding site in Fiji. PLoS one 6, e16597.

[33] Rasalato, E., Maginnity, V. & Brunnschweiler, J.M. 2010 Using local ecological knowledge to identify shark river habitats in Fiji (South Pacific). Environmental Conservation 37, 90–97.

[34] Glaus, K.B., Adrian-Kalchhauser, I., Burkhardt-Holm, P., White, W.T. & Brunnschweiler, J.M. 2015 Characteristics of the shark fisheries of Fiji. Scientific reports 5, 17556.

[35] Glaus, K.B., Appleyard, S.A., Stockwell, B., Brunnschweiler, J.M., Shivji, M., Clua, E., Marie, A.D. & Rico, C. 2020 Insights Into Insular Isolation of the Bull Shark, Carcharhinus leucas (Müller and Henle, 1839), in Fijian Waters. Frontiers in Marine Science 7, 1087.

[36] Brunnschweile, J.M. 2010 The Shark Reef Marine Reserve: A marine tourism project in Fiji involving local communities. Journal of Sustainable Tourism 18. (doi:10.1080/09669580903071987).

[37] Bouveroux, T., Loiseau, N., Barnett, A., Marosi, N.D. & Brunnschweiler, J.M. 2021 Companions and Casual Acquaintances: The Nature of Associations Among Bull Sharks at a Shark Feeding Site in Fiji. Frontiers in Marine Science 8. (doi:10.3389/fmars.2021.678074).

[38] Cardeñosa, D., Glaus, K.B.J. & Brunnschweiler, J.M. 2017 Occurrence of juvenile bull sharks (Carcharhinus leucas) in the Navua River in Fiji. Marine and Freshwater Research 68. (doi:10.1071/MF16005).

[39] Glaus, K.B.J., Brunnschweiler, J.M., Piovano, S., Mescam, G., Genter, F., Fluekiger, P. & Rico, C. 2019 Essential waters: Young bull sharks in Fiji’s largest riverine system. Ecology and Evolution 9. (doi:10.1002/ece3.5304).

[40] Brunnschweiler, J., Marosi, N. & Glaus, K. 2024 Connecting the dots: tracking bull sharks from a provisioning site into the species’ river parturition sites in Fiji. Pacific Conservation Biology 30.

[41] Lemopoulos, A., Prokkola, J.M., Uusi-Heikkilä, S., Vasemägi, A., Huusko, A., Hyvärinen, P., Koljonen, M.L., Koskiniemi, J. & Vainikka, A. 2019 Comparing RADseq and microsatellites for estimating genetic diversity and relatedness — Implications for brown trout conservation. Ecology and Evolution 9, 2106–2120. (doi:10.1002/ece3.4905).

[42] Santure, A.W., Stapley, J., Ball, A.D., Birkhead, T.R., Burke, T. & Slate, J. 2010 On the use of large marker panels to estimate inbreeding and relatedness: Empirical and simulation studies of a pedigreed zebra finch population typed at 771 SNPs. Molecular Ecology 19. (doi:10.1111/j.1365-294X.2010.04554.x).

[43] Duncan, K.M. & Holland, K.N. 2006 Habitat use, growth rates and dispersal patterns of juvenile scalloped hammerhead sharks Sphyrna lewini in a nursery habitat. Marine Ecology Progress Series 312. (doi:10.3354/meps312211).

[44] Kilian, A., Wenzl, P., Huttner, E., Carling, J., Xia, L., Blois, H., Caig, V., Heller-Uszynska, K., Jaccoud, D. & Hopper, C. 2012 Diversity arrays technology: a generic genome profiling technology on open platforms. In Data production and analysis in population genomics: Methods and protocols (pp. 67–89, Springer.

[45] Marie, A.D., Herbinger, C., Fullsack, P. & Rico, C. 2019 First reconstruction of kinship in a scalloped hammerhead shark aggregation reveals the mating patterns and breeding sex ratio. Frontiers in Marine Science 6, 676.

[46] Jones, O.R. & Wang, J. 2010 COLONY: a program for parentage and sibship inference from multilocus genotype data. Molecular ecology resources 10, 551–555.

[47] Gu, Z., Gu, L., Eils, R., Schlesner, M. & Brors, B. 2014 Circlize implements and enhances circular visualization in R. Bioinformatics 30. (doi:10.1093/bioinformatics/btu393).

[48] Wang, J. 2002 An estimator for pairwise relatedness using molecular markers. Genetics 160, 1203–1215. (doi:10.1093/genetics/160.3.1203).

[49] Pew, J., Muir, P.H., Wang, J. & Frasier, T.R. 2015 related: An R package for analysing pairwise relatedness from codominant molecular markers. Molecular Ecology Resources 15. (doi:10.1111/1755-0998.12323).

[50] Csardi, G. & Nepusz, T. 2006 The igraph software. Complex syst 1695, 1–9.

[51] Do, C., Waples, R.S., Peel, D., Macbeth, G.M., Tillett, B.J. & Ovenden, J.R. 2014 NeEstimator v2: Re-implementation of software for the estimation of contemporary effective population size (Ne) from genetic data. Molecular Ecology Resources 14, 209–214. (doi:10.1111/1755-0998.12157).

[52] Klein, J.D., Bester-van der Merwe, A.E., Dicken, M.L., Mmonwa, K.L. & Teske, P.R. 2019 Reproductive philopatry in a coastal shark drives age-related population structure. Marine Biology 166, 26.

[53] Portnoy, D., Puritz, J.B., Hollenbeck, C.M., Gelsleichter, J., Chapman, D. & Gold, J. 2015 Selection and sex biased dispersal in a coastal shark: The influence of philopatry on adaptive variation. Molecular Ecology 24, 5877–5885.

[54] Knip, D.M., Heupel, M.R. & Simpfendorfer, C.A. 2012 To roam or to home: Site fidelity in a tropical coastal shark. Marine Biology 159. (doi:10.1007/s00227-012-1950-5).

[55] Feldheim, K.A., Gruber, S.H. & Ashley, M.V. 2004 Reconstruction of parental microsatellite genotypes reveals female polyandry and philopatry in the lemon shark, Negaprion brevirostris. Evolution 58, 2332–2342.

[56] Acuña-Marrero, D., Smith, A.N., Hammerschlag, N., Hearn, A., Anderson, M.J., Calich, H., Pawley, M.D., Fischer, C. & Salinas-de-León, P. 2017 Residency and movement patterns of an apex predatory shark (Galeocerdo cuvier) at the Galapagos Marine Reserve. PLoS One 12, e0183669.

[57] Paris, A. 2020 Dreketi River and Estuary Shark and Ray Survey. (pp. 1–19. Suva, Fiji, WWF-Pacific.

[58] Brunnschweiler, J.M. & Compagno, L.J. 2008 First record of Carcharhinus leucas from Tonga, South Pacific. Marine Biodiversity Records 1, e51.

[59] Palstra, F.P. & Ruzzante, D.E. 2008 Genetic estimates of contemporary effective population size: what can they tell us about the importance of genetic stochasticity for wild population persistence? Molecular ecology 17, 3428–3447.

[60] Frankham, R., Bradshaw, C.J. & Brook, B.W. 2014 Genetics in conservation management: revised recommendations for the 50/500 rules, Red List criteria and population viability analyses. Biological Conservation 170, 56–63.

[61] Balloux, F. & Lehmann, L. 2003 Random mating with a finite number of matings. Genetics 165, 2313–2315.

[62] Tsai, W.-P., Liu, K.-M., Punt, A.E. & Sun, C.-L. 2015 Assessing the potential biases of ignoring sexual dimorphism and mating mechanism in using a single-sex demographic model: the shortfin mako shark as a case study. ICES Journal of Marine Science 72, 793–803.

[63] Bernal, M., Sinai, N., Rocha, C., Gaither, M., Dunker, F. & Rocha, L. 2015 Long term sperm storage in the brownbanded bamboo shark Chiloscyllium punctatum. Journal of fish biology 86, 1171–1176.

[64] Holt, W. & Lloyd, R. 2010 Sperm storage in the vertebrate female reproductive tract: how does it work so well? Theriogenology 73, 713–722.

[65] Jacoby, D.M.P., Croft, D.P. & Sims, D.W. 2012 Social behaviour in sharks and rays: Analysis, patterns and implications for conservation. Fish and Fisheries 13. (doi:10.1111/j.1467-2979.2011.00436.x).

[66] Papastamatiou, Y.P., Mourier, J., Pouca, C.V., Guttridge, T.L. & Jacoby, D.M.P. 2022 Shark and Ray Social Lives. In Biology of Sharks and Their Relatives (

[67] Dwyer, R.G., Krueck, N.C., Udyawer, V., Heupel, M.R., Chapman, D., Pratt, H.L., Garla, R. & Simpfendorfer, C.A. 2020 Individual and Population Benefits of Marine Reserves for Reef Sharks. Current Biology 30, 480–489.e485. (doi:10.1016/j.cub.2019.12.005).

[68] Gwinn, D.C., Allen, M.S., Johnston, F.D., Brown, P., Todd, C.R. & Arlinghaus, R. 2015 Rethinking length based fisheries regulations: the value of protecting old and large fish with harvest slots. Fish and Fisheries 16, 259–281.

[69] Vignaud, T.M., Meyer, C.G., Séguigne, C., Bierwirth, J. & Clua, E.E.G. 2023 Examining individual behavioural variation in wild adult bull sharks (Carcharhinus leucas) suggests divergent personalities. Behaviour 160. (doi:10.1163/1568539X-bja10244).

